# Molecular Signatures for Microbe-Associated Colorectal Cancers

**DOI:** 10.1101/2024.05.26.595902

**Authors:** Ibrahim M Sayed, Daniella T Vo, Joshua Alcantara, Kaili M Inouye, Rama F Pranadinata, Linda Luo, C Richard Boland, Nidhi P. Goyal, Dennis John Kuo, Sherry C. Huang, Debashis Sahoo, Pradipta Ghosh, Soumita Das

**Affiliations:** Department of Pathology, University of California San Diego, La Jolla, CA, USA; Department of Pediatrics, University of California San Diego, La Jolla, CA, USA; Department of Computer Science and Engineering, Jacob’s School of Engineering, University of California San Diego, La Jolla, CA, USA; Department of Cellular and Molecular Medicine, University of California San Diego, San Diego, CA, USA; Rady Children’s Institute for Genomic Medicine, San Diego, CA, USA; Department of Medicine, University of California San Diego, La Jolla, CA, USA; Department of Pediatrics, Rady Children’s Hospital, San Diego, San Diego, CA, USA; Moores Cancer Center, University of California San Diego, San Diego, CA, USA

**Keywords:** Artificial intelligence, gene signatures, patient-derived organoids, familial adenomatous polyposis (FAP), APC^Min+/-^ mice, colorectal cancer (CRC)-associated microbes, *Fusobacterium nucleatum (Fn)*

## Abstract

**Background:** Genetic factors and microbial imbalances play crucial roles in colorectal cancers (CRCs), yet the impact of infections on cancer initiation remains poorly understood. While bioinformatic approaches offer valuable insights, the rising incidence of CRCs creates a pressing need to precisely identify early CRC events. We constructed a network model to identify continuum states during CRC initiation spanning normal colonic tissue to pre-cancer lesions (adenomatous polyps) and examined the influence of microbes and host genetics.

**Methods:** A Boolean network was built using a publicly available transcriptomic dataset from healthy and adenoma affected patients to identify an invariant Microbe-Associated Colorectal Cancer Signature (MACS). We focused on *Fusobacterium nucleatum* (*Fn*), a CRC-associated microbe, as a model bacterium. MACS-associated genes and proteins were validated by RT-qPCR, RNA seq, ELISA, IF and IHCs in tissues and colon-derived organoids from genetically predisposed mice (*CPC-APC^Min+/-^*) and patients (FAP, Lynch Syndrome, PJS, and JPS).

**Results:** The MACS that is upregulated in adenomas consists of four core genes/proteins: CLDN2/Claudin-2 (leakiness), LGR5/leucine-rich repeat-containing receptor (stemness), CEMIP/cell migration-inducing and hyaluronan-binding protein (epithelial-mesenchymal transition) and IL8/Interleukin-8 (inflammation). MACS was induced upon *Fn* infection, but not in response to infection with other enteric bacteria or probiotics. MACS induction upon *Fn* infection was higher in *CPC-APC*^Min+/-^ organoids compared to WT controls. The degree of MACS expression in the patient-derived organoids (PDOs) generally corresponded with the known lifetime risk of CRCs.

**Conclusions:** Computational prediction followed by validation in the organoid-based disease model identified the early events in CRC initiation. MACS reveals that the CRC-associated microbes induce a greater risk in the genetically predisposed hosts, suggesting its potential use for risk prediction and targeted cancer prevention.

## Introduction

Colorectal carcinoma (CRC) represents the third most prevalent cancer worldwide [1], with the global incidence predicted to increase by 60%, resulting in ∼1.1 million deaths annually, by 2030 [2]. According to the American Cancer Society, about 5% of CRC cases are associated with inherited mutations linked to cancer predisposition syndromes. The most common are Lynch syndrome (hereditary non-polyposis colorectal cancer, or HNPCC) [3] and familial adenomatous polyposis (FAP) [4, 5]. Lynch Syndrome involves the loss of functions of the mismatch repair genes MLH1 (mutL homolog 1), MSH2 (mutS homolog 2), MSH6 (mutS homolog 6), and PMS2 (postmeiotic segregation 2), as well as the gene EPCAM (epithelial cellular adhesion molecule) [6]. Conversely, FAP is characterized by loss of function of the adenomatous polyposis coli (APC) gene which is a negative regulator of WNT/β-Catenin, a proliferative signaling pathway whose upregulation is associated with cancer development. Other conditions like MUTYH-associated polyposis and certain hamartomatous polyposis conditions, including Peutz-Jeghers Syndrome (PJS) and Juvenile Polyposis Syndrome (JPS) are also linked to increased CRC risk [7, 8].

Apart from genetic predispositions and non-modifiable risk factors, approximately 70-75% of CRC cases are associated with modifiable factors such as lifestyle, diet, and gut microbiota [9]. Omics-technology has been utilized to investigate the gut microbiota’s association with cancer initiation [10]. Recent studies have shown that specific commensals can collectively induce CRC by generating DNA damage from the microbe-derived metabolites [11]. Fecal metagenomic and metabolomic studies in large cohorts have demonstrated that shifts in microbial populations and their metabolite profiles are linked to CRC[12]. It has been hypothesized that a compromised gut barrier may allow microbes and toxins to cross the gut epithelium triggering inflammation that fuels CRC initiation and progression [13–16]. Microbial biofilms are implicated in driving CRCs in Lynch and FAP syndromes; genetically engineered mouse models show that specific bacteria can induce colon inflammation and tumor formation [17, 18]. The transplantation of conventional microbiota increased microsatellite instability in untransformed intestinal epithelium of Msh2-Lynch mice, indicating that the microbial composition influences the rate of mutagenesis in MSH2-deficient crypts [19]. Pathogenic bacteria such as enterotoxigenic *Bacteroides fragilis* and *pks+ E. coli* have been linked to colitis-associated CRCs, and *Fusobacterium nucleatum* (*Fn*) has been linked to sporadic CRCs [20–25]. Other CRC-associated microbes include *Clostridium difficile*, *Enterococcus faecalis, Helicobacter pylori, and Streptococcus gallolyticus (formerly known as S. bovis-type I)* [20]. Despite increasing evidence of these associations, how microbes fuel CRCs remains poorly understood [26], and the individual and combined contribution(s) of microbes and/or host genetics towards the risk of CRCs remains to be defined.

To address this gap, we have generated a computational map to identify the gene expression changes during the normal colonic tissue to adenoma progression (**Fig. 1**). We validated these network-derived signatures in genetically predisposed human and mouse colon samples. To assess the gene signatures, we developed patient-derived organoids (PDOs) from the colonic tissues of subjects with hereditary polyposis syndromes. Finally, we used 3D organoid models and 2D enteroid-derived monolayers co-cultured with CRC-associated microbes to understand the impact of the microbes on host gene signatures linked to cancer initiation.

**Figure 1:**
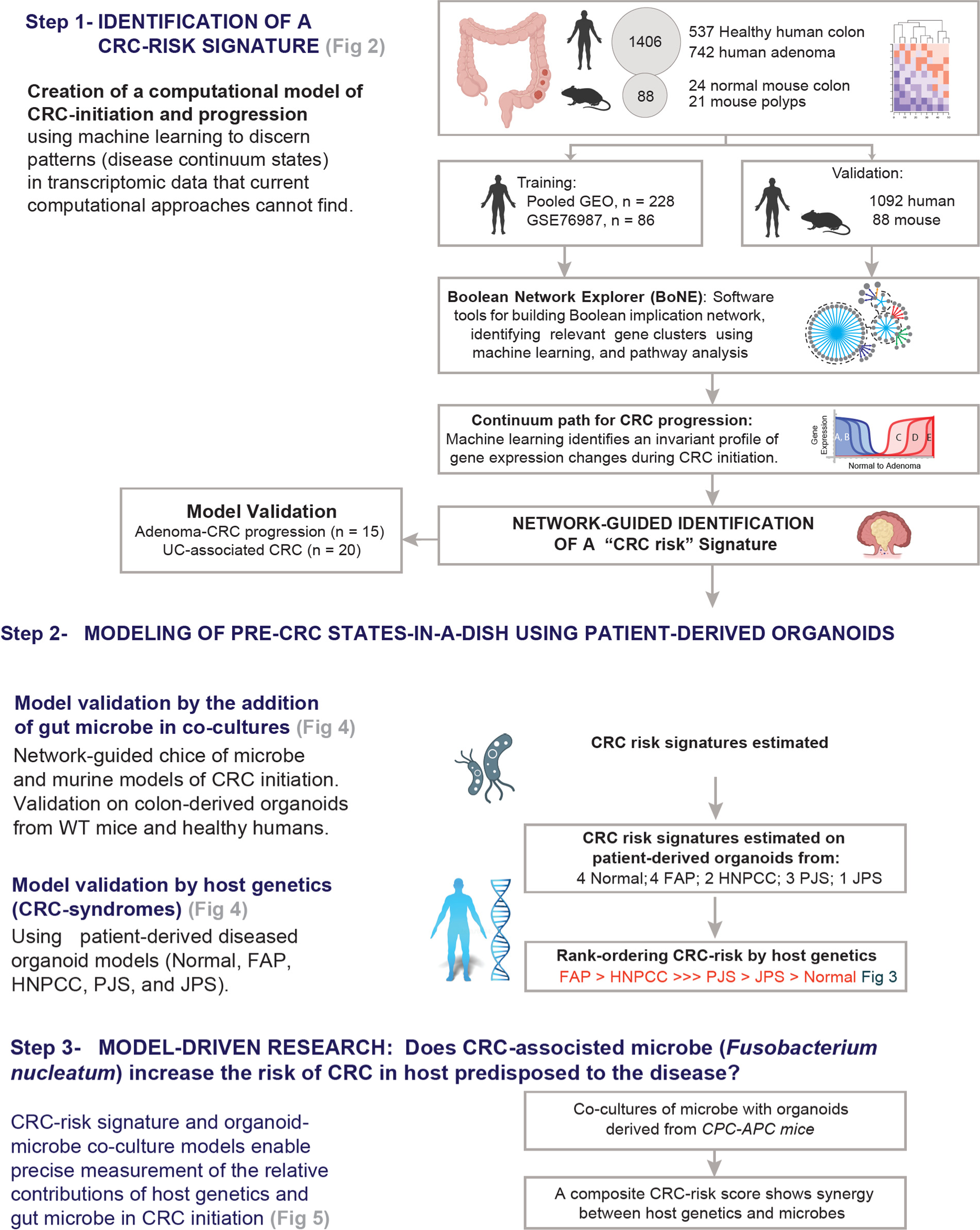
Study design. A database including 1567 gene expression data from both 1406 human samples and 88 mouse samples was used to build up a computational model of colorectal carcinoma (CRC) via a Boolean Network Explorer (BoNE). Boolean implication network using machine learning identifies invariant gene signatures during CRC initiation and progression. These gene signatures were tested in patient-derived diseased organoid models (Normal, FAP, Lynch Syndrome, PJS, and JPS). Moreover, these transcriptome changes were tested in murine models of CRC initiation challenged with CRC-associated microbe (*Fn*) that could drive cancer initiation and/or progression where the presence of host genetic factors further triggers the cancer initiation and progression.

## Results

### Boolean implication network identifies key pathways in the progression from normal colonic tissue to adenoma

Our computational algorithms have enhanced our ability to analyze “big data”, such as transcriptomics, with the goal of understanding complex human diseases and prioritizing diagnostic, prognostic, and therapeutic markers. For example, methods using networks to map relationships between genes have been widely utilized for understanding human diseases [27–32]. Pair-wise gene relationships are usually identified through methods implementing correlation [33–38], mutual information [30], or other linear-based computational techniques [39] dimension reduction [40] and clustering [41, 42]. However, these results may not be reproducible in the real world. A method to build a network using transcriptomics data, in which gene clusters are connected by Boolean invariant relationships [43, 44], has been used to track the progression of cellular states along any disease continuum. Previously, this methodology was used to identify translationally relevant cellular states with high degrees of accuracy in diverse samples and tissues [45–58]. Most recently, it was used to identify a therapeutic target to protect the gut barrier in inflammatory bowel disease [44].

Using publicly available transcriptomic datasets of normal and adenoma colonic tissues (**Fig. 2A, Supplementary Figure S1, Supplementary Table 1)**, a Boolean implication network was built using BoNE [44] (Boolean Network Explorer), where few large clusters were formed, while smaller-sized clusters were the most common (**Fig. 2B)**. The Boolean implication network showed charting of the Boolean paths (**Fig. 2C)** where clusters are linked to each other through the six possible Boolean relationships (**Fig. 2C-2D**). Reactome pathway analysis of these clusters along the path continuum revealed the most important biological processes involved in normal to adenoma progression (**Fig. 2E**). Each cluster was put on the normal or adenoma side depending on the average gene expression value of a cluster, and then the clusters were organized from normal (left) to adenoma (right), showing the biological processes during the initiation and progression of adenoma (**Fig. 2E, Supplementary Table 2)**, as described previously in IBD network [44]. There is a clear initiation cluster (C#1) and an equally clear termination cluster (C#4 and C#5); multiple paths converged on the latter. C#1, which was the farthest cluster in the normal colon, was enriched primarily in pathways that are initiated by AMPK (AMP-activated protein kinase) **(Supplementary Table 2)**. The termination cluster C#4 was enriched in βCatenin/TCF/LEF Wnt signaling programs, their target genes and cell-cell junction organization and C#5 include TLRs, MyD88 involved in microbial sensing **(Supplementary Table 2).** These initiation and termination clusters also represented elements of either genetic predisposition, injury and/or immune activation (Chemokine receptors, MAPK, IL-6 and T-cell response) or oncogene activation (PI3K and KRAS). Common genetic and epigenetic aberrations known to propel adenoma progression were seen: KRAS (C#7), BRAF (C#4 and C#14), NOTCH and Eph/Ephrin signaling (C#6), cell-cycle and DNA damage (C#13, C#15). Interestingly, we also found an IL8-specific signature in adenomas (C#4) (**Fig. 2E**).

**Figure 2:**
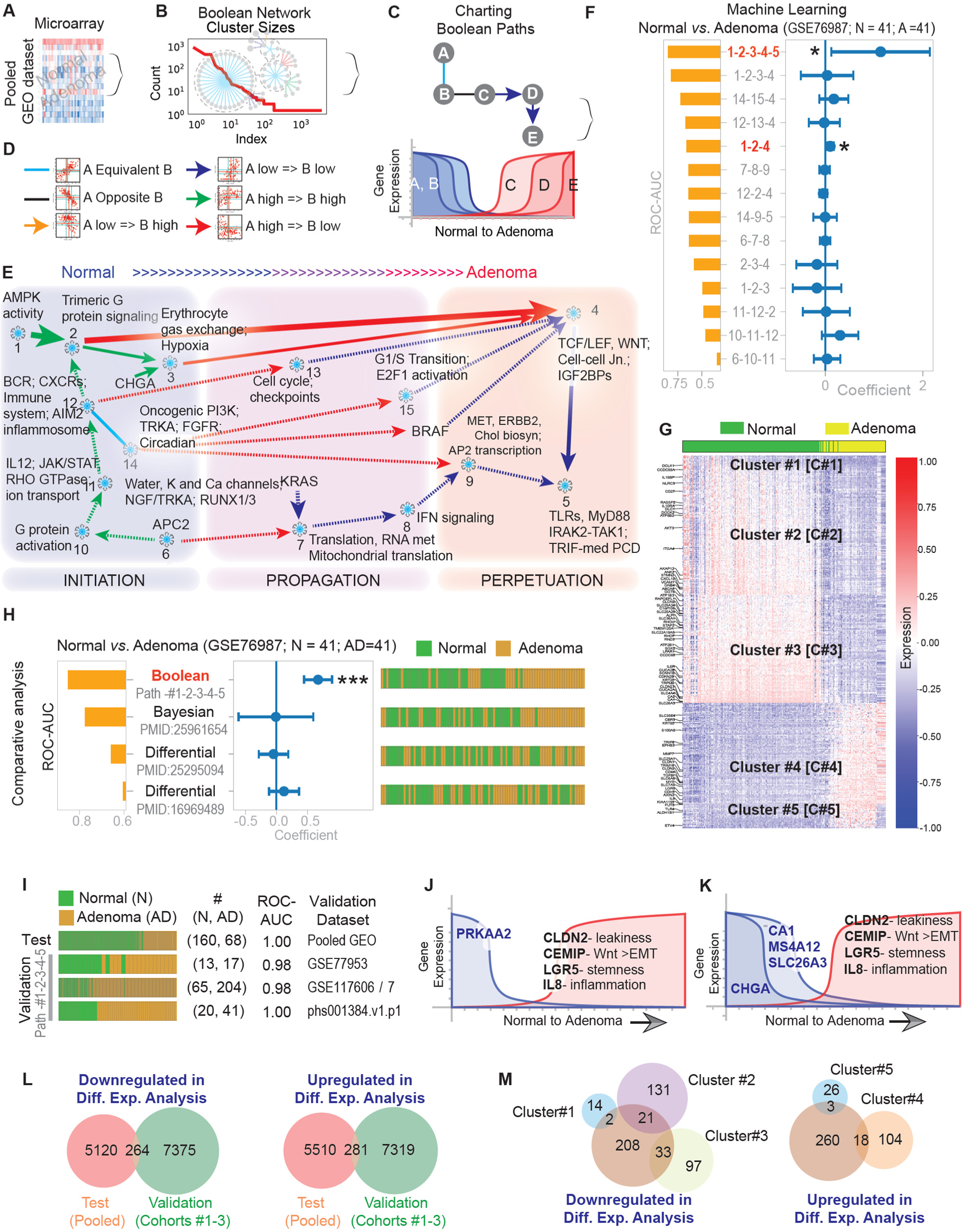
Boolean Network map of Colonic Adenomas; Establishment and Evaluation. **A)** Boolean network analysis was performed on a pooled transcriptomics dataset of normal colon and adenoma dataset downloaded from Gene Expression Omnibus (see Materials and Methods) to identify global pathways and biological processes that are enriched in a continuum of cellular states from normal colon to adenoma. (**B-D**) Genes with similar expression profiles were organized into clusters, and relationships between clusters were represented as color-coded edges connecting clusters. **B)** Graph displaying the distribution of the sizes of clusters of genes that are equivalent to each other confirms scale-free architecture of the Boolean network in A. **C.)** Top: Schematic showing an example of how Boolean cluster relationships are used to chart disease paths, beginning with the largest cluster in healthy controls and following specific Boolean invariants: A low => B low, A opposite B, and A high => B high. Bottom: Schematic showing individual gene expression changes along a Boolean path within the normal to adenoma continuum, illustrating the changing levels of expression of the genes in the above example. Boolean network contains the six possible Boolean relationships between genes as color-coded edges connecting clusters. **E)** Reactome pathway analysis of each cluster along the top continuum paths was performed to identify the signaling pathways and cellular processes that are enriched during adenoma initiation and progression. **F)** Selection of the Boolean path using machine learning. GSE76987 data set was used to select the best path that can separate normal (n=41) and adenoma (n=41) samples. The coefficient of each path score (at the center) with 95% confidence intervals (as error bars) and the p values expressed as significant (*) were illustrated in the bar plot. The p-value for each term tests the null hypothesis that the coefficient is equal to zero (no effect). Clusters C#1-2-3-4-5, and cluster C#1-2-4 showed the best performance. **G)** A heatmap of the expression profile of genes within Boolean clusters superimposed on sample type (top bar) shows the accuracy of Boolean analysis in sample segregation into normal and adenomas. **H)** Comparison of Boolean (using the C#1-2-3-4-5 path), Bayesian, and Differential analysis in segregation Normal from adenoma samples in the dataset GSE76987. **I)** Validation of Boolean analysis (the C#1-2-3-4-5 path) in testing other data sets (Pooled GEO, GSE77953, GSE117606/7, and phs001384.v1.p1). **J-K)** Detailed view of two prominent disease paths identifying downregulation of PRKAA2 (Cluster 1; AMPKα2) and CHGA (Chromogranin A) as two early events in the process; both are accompanied by an up-regulation of LGR5, CEMIP, IL8 and CLDN2 (a leaky TJ protein). **L-M)** Overlap of differentially expressed genes (upregulated or downregulated) between two independent datasets (test cohort and Validation cohorts). The overlap of differently expressed genes is illustrated using two Venn diagrams. The overlap between the genes in different clusters (C#1-2-3-4-5) is presented in (**M**) Downregulated genes that overlap in clusters C#1-2-3 (left), and upregulated genes that overlap in C#4-5 (right).

Next, we used machine learning to identify the best gene clusters (nodes) connected by Boolean implication relationships (edges) in distinguishing normal from adenoma samples (**Fig. 2F)**. Using our training dataset (normal samples: n = 41; adenoma samples: n = 41) and different possible cluster combinations, we identified clusters 1-2-3-4-5 (C#1-2-3-4-5) as the best in separating normal samples from adenoma samples with the highest accuracy (**Fig. 2F-2G**). Then we evaluated the C#1-2-3-4-5 path in different cohorts, test cohorts (n=2) and validation cohorts (n=16; 13 human cohorts and 3 mouse cohorts), and we found that the C#1-2-3-4-5 path performed consistently well across all the independent cohorts **(Supplementary Fig. S2-S6).** To assess the accuracy of predicting normal vs adenoma samples using the C#1-2-3-4-5 path vs. other conventional bioinformatics tools, we compared Boolean analysis directly with differential and Bayesian approaches that were used by others for analysis of gene expression profiles in adenoma and CRC samples [59–61]. Using the training dataset, we found that the C#1-2-3-4-5 path was more accurate than the other two approaches and the only significant approach in segregation normal from adenoma samples (**Fig. 2H**). The C#1-2-3-4-5 path could classify normal from adenoma in other validation cohorts with high accuracy (ROC-AUC 0.98-1.00) (**Fig. 2I**). All the previous findings show the power of Boolean networks in accurately modeling gene expression changes that occur during normal to adenoma progression.

Using the disease map in **Fig. 2E**, we found that the catalytic α-2 subunit of AMPK (encoded by the *PRKAA2* gene) was one of the first genes to be downregulated within the most prominent disease path identified (C#1), which was associated with upregulation of 4 key genes in C#4; *CEMIP*; i.e. *KIAA1199*, which encodes the Cell migration-inducing and hyaluronan-binding protein; *CLDN2*, which encodes the cation-selective channels in the paracellular space; *LGR5*, which encodes the stemness-reporter Leucine-rich repeat-containing G-protein coupled receptor, and interleukin-8 (IL8)-predominant inflammation (**Fig. 2J**). Another path with the same endpoint was the downregulation of CHGA (**Fig. 2K),** which was followed by the downregulation of 3 other genes that are often used as markers of differentiation in the colon: Carbonic anhydrase-1 (CA1), Membrane Spanning 4-Domains, Subfamily A, Member 12 (MS4A12), and Solute Carrier Family 26 Member 3 (SLC26A3, i.e. down-regulated in adenoma, DRA) (**Fig. 2K**). The downregulation of CHGA was also associated with the upregulation of C#4 genes mentioned in **Fig. 2J and Fig. 2K**. Differential expression analysis revealed limited overlap between the upregulated and downregulated genes in the test pooled cohort and the three validation cohorts (**Fig. 2L**). The C#1-2-3-4-5 path exhibited minimal overlap in its commonly differentially expressed genes (upregulated and downregulated) (**Fig. 2M**).

### Validation of the invariant gene signatures using human CRC tissues and organoids

Looking at the major genes that are known to be mutated in hereditary CRC, we noticed that many of these genes are not present in C#1-2-3-4-5 but are present in C#6 (APC, STK1), and C#7 (MSH2 and SMAD), which are connected to C#1-2-3-4-5 (**Fig. 3A, Supplementary Table 2).** We validated these predictions by RT-qPCR on colon tissues collected from healthy controls and from non-involved (NI) and polyp tissues collected from the colons of genetically predisposed subjects. We found that the genes along the C#1-2-3-4 path were up/downregulated exactly as predicted: *PRKAA2* and *CHGA* were downregulated concomitantly with the upregulation of *CEMIP*, *CLDN2*, *LGR5* and *IL8* in the predisposed mucosa and in polyps (**Fig. 3B, Fig S7)**. Next, we asked if the C#1-2-3-4-5 path can classify the CRC syndromes based on the established lifetime CRC risk profiles associated with these syndromes, which we gathered from publicly available information related to disease epidemiology, and it is higher in FAP > Lynch > JPS ∼ PJS (**Fig. 3C, Supplementary Table 3).** Next, we performed RNA-seq from genetically predisposed human colon samples and found that the C#1-2-3-4-5 path, used as a gene signature, could separate different samples either derived from non-involved (**Fig. 3D-left, Fig S8)** or polyp region (**Fig. 3D-right, Fig S8)** that are associated with low vs high risk of CRC.

**Figure 3:**
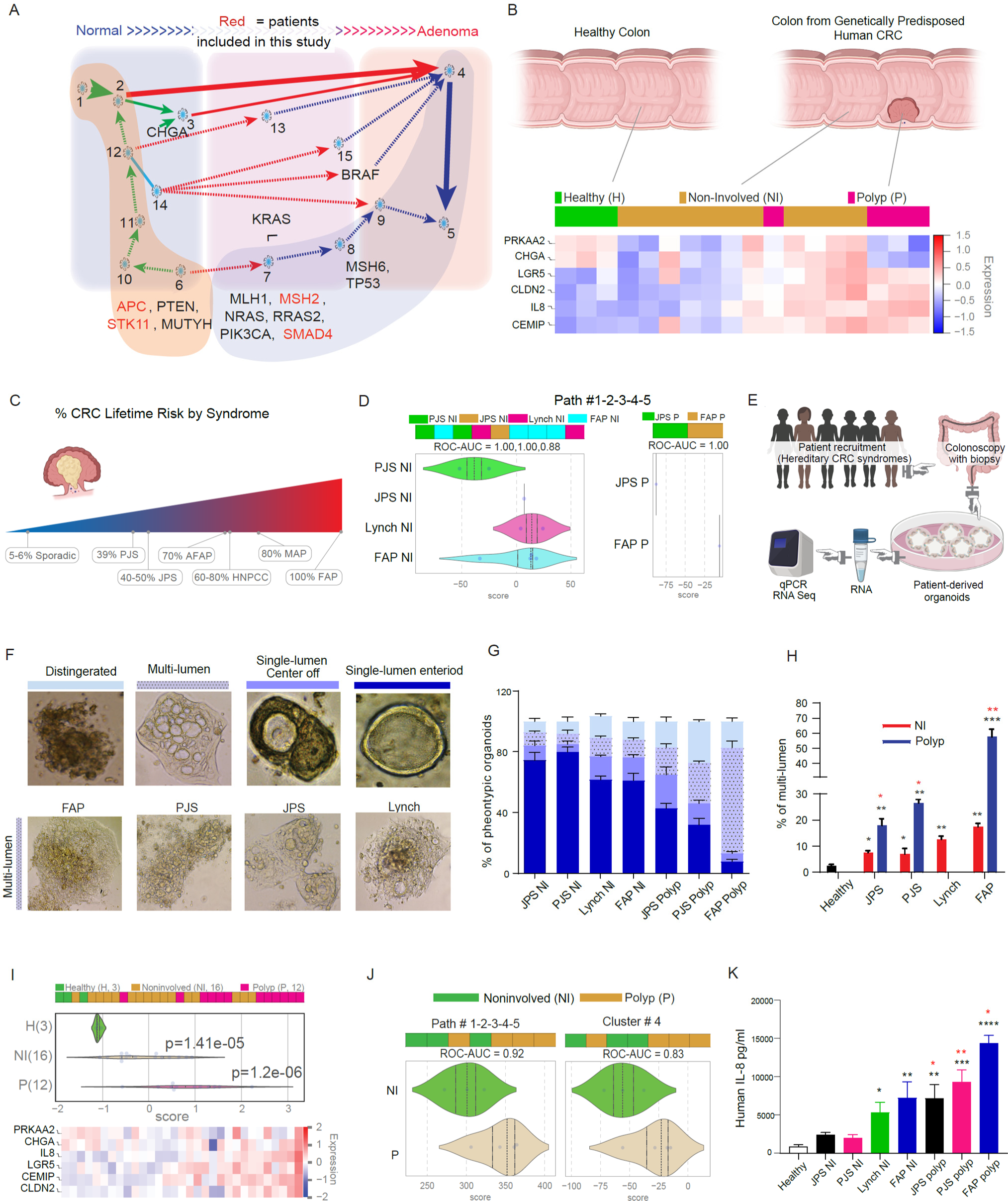
Validation of the invariant gene signatures using human CRC tissues and organoids. **A)** Disease pathways identify the clusters and gene changes during normal to adenoma initiation. **B)** Heatmap of the RT-qPCR gene expression predicted from BoNE using colon tissues derived from healthy human (n=3), and genetically predisposed CRC patients (FAP, Lynch, PJS, and JPS) where non-involved (NI, n= 11) and polyp (P=4) tissues were assessed. See Supplementary Figure 7 for more analysis. Key genes in clusters 1,3, and 4 are presented on the left of the heatmap. Data presented as DCT (Ct of target gene-Ct of the endogenous control gene). The expression level ranged from -1 to +1; the expression below 0 indicated low expression and represented with blue color and the expression above 0 indicated high expression and represented with red color. **C)** Schematic showing the risk of CRC in different hereditary CRC syndromes, where FAP (red) represents the highest risk of CRC development and sporadic (blue) represents the lowest risk and other categories in between them. Data collected from the published literature and described in **Supplementary Table 3**. **D)** Bar and violin plots from non-involved tissues (left) and polyp tissues (right). Violin plots display the rank ordering of different CRC patient samples using the average gene expression patterns of the invariant genes analyzed by RNA sequencing. ROC-AUC statistics are measured to determine the classification strength of the samples. Schematic showing the study design involving the development of 3D organoids from hereditary polyposis samples and assessment of the invariant gene signatures using RT-qPCR and RNA seq. **F)** Major phenotypic features observed in organoid lines derived from hereditary CRC syndromes. Top: 4 morphological phenotypic organoids were presented (disintegrated (light blue), multi-lumen organoids (dotted violet), single lumen off-center (azure), and single lumen (dark blue)). Bottom: representatives of multi-lumen organoids from different hereditary CRC syndromes. **G)** The percentage of each organoid phenotype (as color coded in Fig. 3F) in different CRC background were presented. **H)** The percentage of multi-lumen organoids in organoids derived from the colon of healthy human (black), colon of non-involved region from genetic hereditary CRC patients (red), and colon of polyp region from genetic hereditary CRC patients (blue). * represents the difference between CRC and healthy, and * represents the difference between polyp and NI from the same genetic background group. **I)** Gene expression analysis of patient-derived organoids targeting the invariant gene signatures using qRT-PCR. Top: represent bar and violin plots show the rank ordering of organoid samples (using *PRKAA2*, *CHGA*, *CLDN2*, *CEMIP*, *IL8* and *LGR5*) derived from healthy individual’s vs organoid-derived from non-involved regions vs organoid-derived from polyp regions of hereditary CRC syndromes (Samples used: 4 FAP, 2 Lynch, 2 PJS and 1 JPS PDOs isolated from 2 different locations. Each experiment has at least two technical repeats); Bottom: heat map displays the target genes of analysis. **J)** Bar and violin plot showing the expression of genes in the C#1-2-3-4-5 path (left) or C #4 only (right) in RNA seq data from non-involved (NI from 1 FAP and 2 Lynch) and polyp (P from 3 FAP and 1 PJS) regions. **K)** The level of IL-8 cytokine was measured in the supernatant of 3D organoids derived from healthy and different genetic CRC patients by ELISA. * Represents the difference in the level IL8 between CRC (either NI or P) and healthy, and * represents the difference between polyp and NI from the same genetic background group. The same color represents NI and polyp derived from the same genetic background. *, **, *** means p < 0.05, 0.01, and 0.001, respectively as determined by Student’s t-test.

To determine if the colonic epithelium shows network-predicted gene expression changes, we isolated the stem cells from the colonic tissues of genetically predisposed CRC patients and healthy individuals, then grew them as 3D organoid models, and then performed transcriptome analysis on the culturing organoids (**Fig. 3E**). Four distinct phenotypical features were observed during the process of their derivation: a) disintegrated (loosely packed) organoids, b) multi-lumen organoids, c) organoids with a single, but center-off lumen, and d) organoids with a centrally located lumen (**Fig. 3F *top row*, samples listed in Supplementary Table 4).** The multi-lumen organoids are collections of several organoids that appeared to be connected (**Fig. 3F *bottom row*).** We hypothesized the multi-lumen structures could be related to epithelial-mesenchymal transition (EMT). Multi-lumen structures were commonly encountered and were seen at a higher frequency when organoids were derived from polyp (P) regions compared to organoids derived from non-involved (NI) regions of the same patient in all the CRC categories. The abundance of these multi-lumen structures correlated with the lifetime risk of CRC development, i.e., the percentage of multi-lumen organoids is in the following direction: FAP > Lynch > PJS ∼ JPS (**Fig. 3H**). Analysis of the transcriptome of the PDOs by qRT-PCR showed the predicted gene expression profile from Boolean analysis (**Fig. 3I**). RNA-seq data derived from these organoids revealed that the C#1-2-3-4-5 path could segregate organoids derived from the NI region from organoids derived from the polyp region with high accuracy in CRC syndromes (**Fig. 3J, Supplementary Fig. S7-S9).**

Next, we assessed the level of C#4 genes in the protein level, especially IL-8, CEMIP, and CLDN2. Measuring the level of IL-8 cytokines in the culture of PDOs, we found the level of IL-8 was significantly higher in the supernatant of cultured organoids derived from polyp regions compared to the NI regions of the same patients in all CRC categories (**Fig. 3K),** and the level of IL-8 was matched with the lifetime CRC development (**Fig. 3K, Supplementary Table 3).** In addition, immunofluorescence (IF) analyses showed the expression of Claudin-2 was higher in organoids derived from FAP patients compared to organoids derived from healthy individuals, and the expression was much higher in organoids derived from polyp regions compared to organoids from NI regions (**Fig. 4A**). In a complementary approach, we used IHC staining, and it revealed that CEMIP staining was significantly higher in FAP-derived organoids than organoids derived from healthy subjects (**Fig. 4B**).

**Figure 4:**
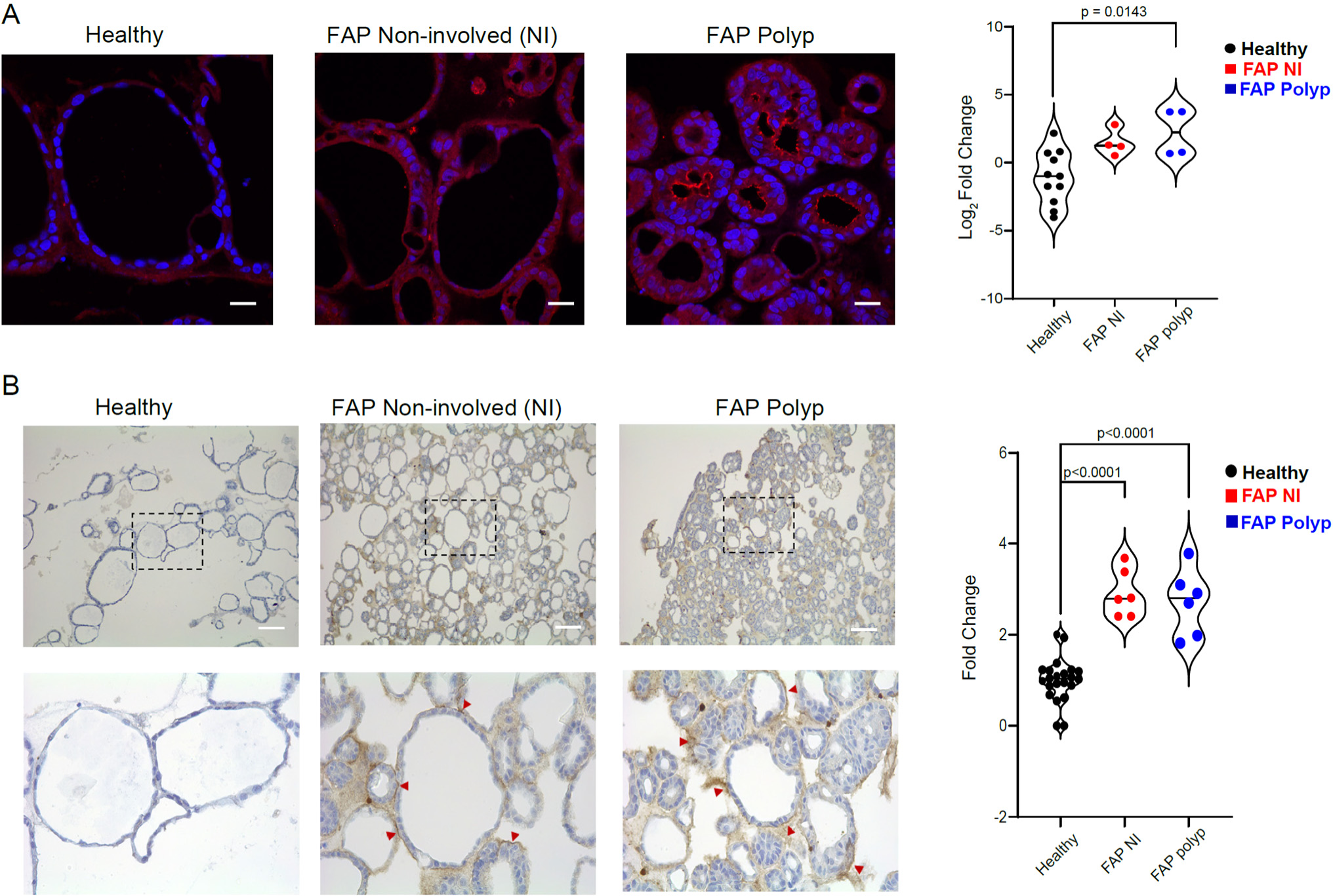
Validation of MACS proteins, CLDN2 and CEMIP by IF and IHC. **A)** Expression of CLDN2 was tested in healthy (black, grey, n=4), FAP NI (red, n=2) and FAP polyp (blue, n=2) PDOs by IF. **(left)** representative images for each PDOs group. **(right)** represent log_2_ fold change expression of CLDN2 intensity signals of FAP divided by the intensity of CLDN2 in healthy organoids. **B)** Expression of CEMIP was tested in healthy (black, grey, n=4), FAP NI (red, n=2) and FAP polyp (blue, n=2) PDOs by IHC. **(left)** representative images for each PDOs group. **(right)** represents fold change expression of CEMIP. **Fold change** = **FAP** [(No. CEMIP positive organoids/ the total number of counted organoids)/ **Healthy** (No. CEMIP positive organoids/ the total number of counted organoids)]

### Cancer-adjacent polyps and colon cancer-associated microbes recapitulate the gene expression changes observed during CRC initiation and progression

We asked if the changes in gene expression along the C#1-2-3-4-5 path are associated with the risk of polyp to CRC progression. To this end, we leveraged a publicly available dataset that represents a time-lapse model for CRC initiation and progression in humans [62]. In that model, cancer adjacent polyps (CAPs) were used as a model to study cancer progression temporally because the precursor polyp of origin remains in direct contiguity with its related polyps [63–65]. Cancer-Free Polyp (CFP) cases, on the other hand, are polyps that have remained cancer free, despite sharing similar size, histologic features and degrees of dysplasia as CAPs (**Fig. 5A**). Thus, in this model the laser-dissected pre-neoplastic tissues from the CAPs represent polyps with a proven high risk of CRCs, CFPs represent polyps at low risk and paired normal colons sampled ∼8 cm away from the polyps served as controls. Performance of the C#1-2-3-4-5 path (derived from the Boolean implication network) was better than other methods of analysis, such as Bayesian and differential analysis, as C#4 alone was sufficient to distinguish normal colon samples from adenomas with high accuracy (**Fig. 5B-C**). Furthermore, Boolean analysis using C#4 can separate quiescent UC samples (low risk of CRC) from UC with remote neoplasia (high risk of CRC) (**Fig. 5D**).

**Figure 5:**
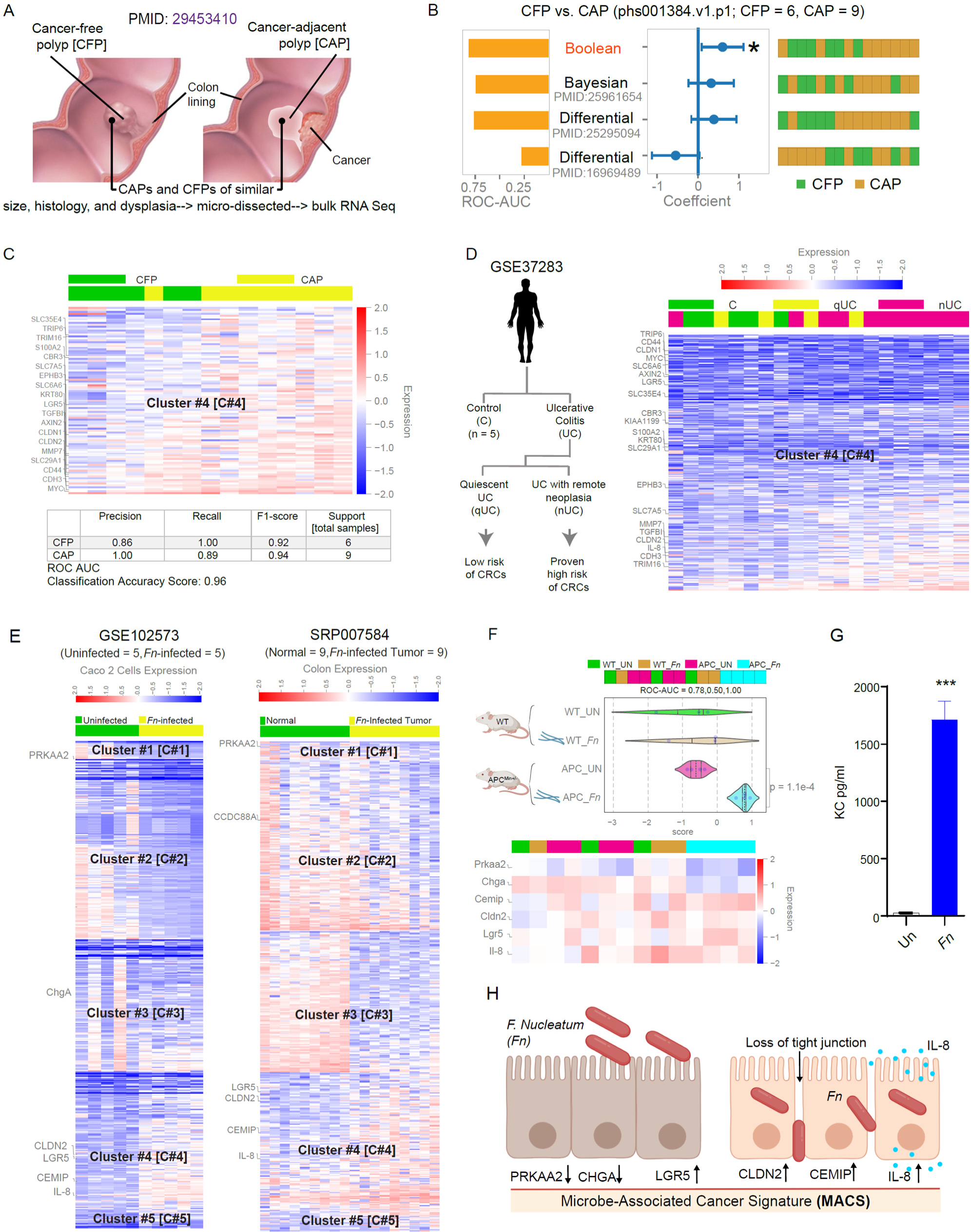
MACS signature detects the risk of CRC conferred by cancer-associated microbes and host genetics. **A)** Schematic showing the previously published (Druliner et al., 2018, PMID:29453410) time-lapse model for CRC initiation and progression. Tubular or villous adenomas that were adjacent to (Cancer-adjacent polyps; CAPs) or free of (Cancer-free polyps; CFPs) cancer and their matched normal colon from each was processed for RNA-Seq. **B)** Comparison of Boolean (the C#1-2-3-4-5 path), Bayesian, and Differential analysis in separating CFP from CAP samples in PMID:29453410. **C)** Bar plot and heatmap of genes in C#4 show the separation of CAP vs CFP samples. **D)** Left: Schematic of study design involving patients with ulcerative colitis-associated CRCs. Right: Bar plot and heatmap of genes in C#4 separates patients with proven risk for CRCs (UC with remote neoplasia; nUC) from those with low risk (quiescent UC; qUC). **E)** Bar plots and heatmaps of genes in the C#1-2-3-4-5 path show the accurate separation of samples into *Fn*-infected or uninfected Caco2 monolayers (left) and *Fn*-infected tumors vs. adjacent normal colon tissues (right). **F)** The gene expression of *Prkaa2*, *Chga*, *Cldn2*, *Cemip*, *Il-8* and *Lgr5* was compared in EDMs developed from WT mouse vs EDMs developed from genetically predisposed APC min mouse, both EDMs were challenged with *Fn*. qRT-PCR was done for *Prkaa2*, *ChgA*, *Cldn2*, *Cemip*, *Il-8* and *Lgr5* and presented as DCT. Top: The bar plot and violin plot show the sample ranking in WT EDMs vs. APC min EDMs that are infected or not with *Fn*. Bottom: Heat map showing the expression of these genes. The expression below 0 indicated low expression and was represented by blue, and the expression above 0 indicated high expression and was represented by red. **G)** The level of Cxcl1 KC (Il-8 homologue) was measured in the supernatant of the EDMs from APC min mice (uninfected and *Fn* infected) as used in (**F**) by ELISA, and the level of cytokine was compared in infected EDMs vs uninfected EDMs. *, *** means p < 0.05 and 0.001 as determined by Student’s t-test. **H)** Summary and proposed working model for the initiation of CRCs by *Fn* through induction of MACS genes. Microbial dysbiosis-mediated stress-induced TJ-collapse and loss of cell polarity which is accompanied by a gene expression signature that is permissive of key cellular processes such as EMT, stemness, leakiness of the gut barrier, and a distinct type of inflammation that is IL-8-predominant. This signature is distinctly associated with dysplastic progression in the epithelium (as in adenomas) and colon cancer initiation within adenomas.

As our main goal is to understand the impact of microbes in the progression of CRC, we searched the publicly available dataset with microbes involved in colorectal carcinogenesis. We analyzed the RNA seq data from organoids and polarized monolayers derived from primary murine colon epithelial cells infected with genotoxic colibactin-producing pks+ *Escherichia coli* strains (**Supplementary Fig. S10)**. Using the C#1-2-3-4-5 path, enteroid-derived monolayers (EDMs) infected with colon cancer microbes can be segregated from non-infected EDMs with high accuracy **(Supplementary Fig. S10).** In contrast, the C#1-2-3-4-5 path cannot separate uninfected compared to infected using probiotics and enteric pathogens **(Supplementary Fig. S11).**

Next, we asked if colorectal cancer-associated microbes can increase the gene expression signature associated with CRC progression. We used the CRC-associated microbe, *Fn* [23] as a model bacteria as it is the most abundant microbe enriched in colonic adenomas and CRCs compared to adjacent normal tissues [23, 66]. *Fn* can trigger tumorigenesis in the *APC*^Min^ mice [23] that resembles human polyposis and is associated with poor survival and chemoresistance among patients with CRCs [67]. Our choice of *Fn* as a model carcinopathogen was further rationalized by the analysis of publicly available transcriptomic datasets with *Fn*-infected colonic epithelial cell lines and *Fn*-associated tumors and adjacent normal tissues [68]. We validated our Boolean prediction using these two publicly available transcriptomic datasets: (a) Caco2 monolayers infected or not with *Fusobacterium nucleatum* (*Fn*) (**Fig. 5E; left**) and (b) *Fn*-infected tumors and adjacent normal tissues (**Fig. 5E; right)**. In both cases, the invariant gene expression signature predicted by the Boolean analysis and that is associated with normal-to-adenoma conversion in the colon could distinguish between *Fn*-infected Caco2 monolayers and tumors from their respective controls (**Fig. 5E**). Specifically, the 6 genes (*Prkaa2*, *Chga*, *Cemip*, *Cldn2*, *Lgr5* and *Il-8*) that are differentially expressed in the NI and polyp area also distinguish between uninfected and *Fn*-infected Caco2 monolayers. We classified these 6 genes as microbe-associated cancer signature (**MACS**). To understand the impact of CRC-associated *Fn* in genetically predisposed conditions, we used *Fn* in the genetically CRC-predisposed murine model. We used CPC-APC^Min/+^ mice strains and their WT littermates. APC^Min/+^ mice have a point mutation in the *Apc* gene and are a model for human FAP. We chose the CPC-APC^Min/+^ mice [69], which carry a CDX2P-NLS Cre recombinase transgene and a flox-targeted Apc allele to achieve a colon-specific gene depletion. Much like human CRCs, these mice develop tumors predominantly in the distal colon and display biallelic Apc inactivation, β-catenin dysregulation, global DNA hypomethylation, and aneuploidy.

To assess the effect of genetic background on the initiation and/or progression of CRC in the presence of microbe, we infected WT murine EDMs and CPC-APC^Min/+^ derived EDMs with *Fn* and evaluated the impact of infection on the previously identified MACS genes. We found that *Fn* infection caused upregulation of *Cldn2* and *Il-8* in both the normal (WT) (and the genetically predisposed (CPC-APC^Min/+^-derived) EDMs (**Fig. 5F),** but *Fn*-infection, induced *Cemip* and *Lgr5* in the CPC-APC^Min/+^-derived EDMs, indicating that infection may fuel EMT and stemness only in the genetically predisposed epithelium after infection (**Fig. 5F**).

Because IL-8 was part of the identified Boolean network path (C#4), we assessed the level of IL-8 using ELISA (**Fig. 5G)** and found that *Fn* induces IL-8 production. Collectively, these findings confirm that recapitulating the distinct IL-8-dominant nature of inflammation was identified earlier by Boolean analysis (**Fig. 2**). We concluded that infection with colon cancer-associated microbes (such as *Fn*) triggered the distinct gene expression signature in EDMs that mirrors the changes observed along the normal-to-adenoma disease paths in the human colon. We also conclude that the presence of genetic predisposition could fuel the initiation and/or progression of CRC, especially in the presence of CRC-associated microbe *Fn* (**Fig. 5H**).

## Discussion

This study is inspired by the urgent need to predict the risk for progression from normal colonic tissue to adenomas to CRC. Microbial infections contribute to 20% of all cancers [70–72]. Chronic microbial infections can trigger abnormal immune responses, leading to inflammation, tissue damage, and DNA alterations. Although it is widely believed that infection-induced inflammation predisposes individuals to CRC, the exact mechanisms by which microbes contribute to this risk remain elusive. Compounding the importance of this issue, current therapies for advanced CRCs often fall short in providing a cure. Thus, detection of CRC risk-associated polyps remains an unmet need. Current post-polypectomy guidelines recommend a repeat colonoscopy in 3 to 10 years based on the size, number, and histology of polyps; however, there is no molecular basis for risk assessment for CRC progression in patients who develop these polyps. Additionally, the guidelines for the appropriate interval between surveillance colonoscopies are poorly supported by evidence [73–75]. Our computational approaches, with the predicted gene signature and validation in patient-derived organoids together with colonoscopy screening, could help detect and prevent CRC at an earlier stage. We used tissues and patient-derived organoids from genetically predisposed human CRCs and animal models (CPC-APC models that mimic human FAP samples) to identify the transcriptome changes during CRC initiation and/or progression. In addition, we challenged differentiated organoid-derived colonic cells with CRC-associated microbe-*Fn* to assess the impact of the infection on CRC development in the genetically predisposed model (CPC-APC EDMs).

Artificial intelligence (AI) has been used in the diagnosis and treatment of CRC [76]. AI is also used for the detection of microsatellite instability in CRC [77], accurate recognition of CRC using semi-supervised deep learning on pathological images [78], development of lncRNA signature for improving outcomes in CRC [79], and determination of T1 CRC metastasis risk to lymph nodes [80]. To our knowledge, no study has utilized computational approaches to predict a microbial signature associated with CRC risk that could be useful in preventing cancer initiation. Here, we have used the Boolean implication network to identify the gene signature during adenoma initiation and progression regardless of the phenotypic (histologic) and genotypic heterogeneity within the analyzed dataset. We assessed the predictive capacity of Boolean analysis in the time-lapse model, where CAPs, CFP and their matched normal colons[62] were analyzed [62]. In that model, CAPs were used to study cancer progression temporally since the precursor polyp of origin remains in direct contiguity with its related CRC [63–65]. CFPs are polyps that have remained cancer-free despite the same size, histologic features, and degrees of dysplasia as CAPs. MACS successfully distinguished normal colon from adenomas, indicating that the predicted gene clusters along the major path (C#1-2-3-4-5) identified previously in our NCBI GEO discovery cohort is also valid in this time-lapse cohort. Our analysis showed that C#4 alone was best at segregating CAPs. CAP and colon cancer-associated microbe *Fn* recapitulate the gene expression changes (by C#4 with 122 genes) in cancer tissue and colon cancer cells. RNA-based panels like the 70-gene ‘MammaPrint’ in breast cancer [81, 82] and 18-gene ‘ColoPrint’ in CRC [83–85] were previously reported. Here we further validated RNA-based signature using 6 genes (PRKAA2 from C#1; CHGA connected to C#3, CEMIP, CLDN2, LGR5 and IL8 from C#4) to predict the risk of progression to CRC.

The MACS genes showed that the catalytic α-2 subunit of AMPK (*PRKAA2*) was one of the first genes to be downregulated and the downregulation of PRKAA2 is predicted to have four associated changes: 1) trigger the upregulation of key genes in C#4 that are known to trigger epithelial-mesenchymal transition (EMT) during CRC initiation and progression (*CEMIP*; i.e. KIAA1199, which encodes the Cell migration-inducing and hyaluronan-binding protein [86–88]); 2) promote leakiness in an inflamed gut barrier (*CLDN2,* which encodes the cation-selective channels in the paracellular space, Claudin-2 which is associated with barrier dysfunction [89–92]); 3) induce stemness (*LGR5*, which encodes the stemness-reporter Leucine-rich repeat-containing G-protein coupled receptor 5 [93]); 4) induce interleukin-8 (*IL8*)-predominant inflammation (CC#4). While the expression of CLDN2 has been shown to increase in CRCs [94, 95] and promote stemness [96], the pro-inflammatory cytokine IL-8 has previously been implicated in the induction of proliferation, migration, angiogenesis, stemness and metastatic potential in CRCs [97–100].

To assess the impact of microbes on the initiation and/or progression of CRC, we used a stem-cell-based gut-in-a-dish model co-cultured with CRC-associated microbe *Fn*. We found that *Fn* increases the risk in genetically CRC-predisposed conditions. We found that *Fn*-infection led to the downregulation of *Prkaa2* and *ChgA* and upregulation of *Cldn2* and *Il-8* in both the normal (WT) and the genetically predisposed (CPC-APC or CDX2Cre-APCMin-derived) EDMs; however, *Fn* infection induced *CEMIP* and *Lgr5* exclusively in the CDX2Cre - APC^Min^-derived EDMs, indicating that infection may fuel EMT and stemness only in the genetically predisposed epithelium. *Fn* also recapitulates the distinct Il-8-dominant nature of inflammation identified earlier by Boolean analysis. Similarly, previous studies showed the co-occurrence of Il-8 overexpression with *Fn*-enriched tumors [15], and that *Fn* invasion of CRCs is associated with an Il-8-predominant inflammatory signature [101]. It is noteworthy that although there is continued skepticism regarding if *Fusobacterium* species are passengers [25, 102] (i.e., commensal biofilms from the oral cavity that innocently associate with abnormal colonic crypts in patients with CRCs) or drivers, our studies in organoids provide evidence to support the latter.

This genetic predisposition to CRCs arises from the germline mutations or epimutations in the DNA mismatch repair (MMR) genes MLH1, MSH2, MSH6 and PMS2 in Lynch syndrome, and in APC and MUTYH (recessive inheritance for FAP). Mutations in SMAD4, BMPR1A, STK11 and PTEN predispose to hamartomatous polyps in less common syndromes such as, PJS and JPS. We have tested patient samples from these 4 groups (FAP, Lynch, PJS and JPS) and isolated organoids to validate the expression of MACS genes. This study serves as a foundation for future investigations incorporating more tissue samples and patient-derived organoids (PDOs) with fibroblasts and immune cells to better mimic the microenvironment. Notably, a recent study revealed the presence of the fungus *Candida albicans* in 35 different cancer tissues after translocation from the gut to other organs [103]. Exploring the expression status of MACS after infection with other microbes, including viruses and fungi, presents another promising avenue for research. Changes in genomic integrity, oncogenic signaling, cellular migration, inflammatory states, and epigenetic alterations induced by other microbes require further mechanistic studies.

In conclusion, our findings contribute to a better understanding and prediction of infection-inflammation-driven oncogenesis. Identifying host gene signatures can aid in predicting the risk of CRC in early stages. Since microbes are implicated in other cancers (e.g., pancreas, breast), our findings extend beyond CRCs and may uncover new microbial triggers for cancer initiation.

## Declaration

### Ethics approval and consent to participate

All human specimens are collected from subjects after signing the consent forms following the guidelines of IRB as discussed in the method section.

### Consent for publication

All authors declared permission for publication.

### Availability of data and materials

All RNA seq data is available. All the codes have been released on github: https://github.com/sahoo00/BoNE/blob/master/MACS/weight-optimization.ipynb. Other data and materials will be supplied upon request of the readers.

## Authors’ contributions

Study concept and design: IMS, DJK, SCH, DS, PG, SD Acquisition of data: IMS, DV, JA, KM, DS

Human data resources and logistical supplies: RFP, LL, NPG, DJK, SCH Analysis and interpretation of data; IMS, DV, DS, PG, SD

Drafting of the manuscript: IMS, PG, DS, SD

Critical revision of the manuscript for important intellectual content: SD revised the manuscript after getting comments from all authors. SD and DJK coordinated with all the co-authors. IMS, DV, JA, KMI, RFP, LL, SCH, CRB, NPG, DS, PG provided the input.

Statistical analysis: IMS, DV, DS, SD Obtained funding: DS, PG, SD Study supervision: DS, PG, SD

## Acknowledgements

We are grateful to Courtney Tindle, Dominic Stec, Mackenzie R. Fuller, Ayden G. Fonseca at the UC San Diego HUMANOID™ Center of Research Excellence; Kathryn Bouic at Rady Children’s Hospital; Dr. Stella Rita Ibeawuchi, Leanne Dugan and Dibo Mukhopadhyay at the Das lab for technical and logistical support. We thank Dr. Stefania Tocci for providing suggestions during the final editing of the manuscript. Several figures present in the manuscript were designed using www.biorender.com.

## Key Resources Table

| REAGENT or RESOURCE | SOURCE | IDENTIFIER |
| --- | --- | --- |
| **Antibodies** |  |  |
| Rabbit polyclonal anti-Claudin 2 antibody | Abcam | Cat# ab53032 |
| Rabbit polyclonal anti-KIAA1199 antibody | Proteintech | Cat #: 21129-1-AP |
| Goat anti-rabbit Alexa 594 | Invitrogen | A11012 |
| **Bacterial and virus strains** |  |  |
| *Fusobacterium nucleatum* | ATCC | ATCC-25586 |
| **Biological samples** |  |  |
| Human colonic biopsies | Rady Children hospital, San Diego<br>HUMANOID CoRE, UCSD | Project ID#130794<br>IRB 190105 |
| CPC-APC<sup>Min/+</sup> (Adenomatous polyposis coli) mice colon biopsies | UCSD, Das Lab | Das Lab, protocol # S18086 |
| C567BL/6 mice colon biopsies | UCSD, Das Lab | Das Lab, protocol # S18086 |
| **Chemicals, peptides, and recombinant proteins** |  |  |
| Quick-RNA Microprep kit | Zymo Research | Cat# R1050 |
| Direct-zol<sup>TM</sup> RNA MiniPrep | Zymo Research | Cat# R2051 |
| qScript<sup>TM</sup> cDNA SuperMix | Quantabio | Supplier # 95048-100 |
| 2× SYBR Green qPCR Master Mix | Biomake | Cat#: B21703 |
| Human organoid media | HUMANOID CoRE, UCSD | Details of media composition [104] |
| Mouse organoid media | HUMANOID CoRE, UCSD | Details of media composition [104] |
| 5% EDM differentiation media | HUMANOID CoRE, UCSD | Details of media composition [104] |
| Corning® Matrigel® Basement Membrane Matrix, LDEV-free, 10 mL | CORNING | Cat# 354234 |
| TrypLE<sup>TM</sup> Select Enzyme (1X), no phenol red | ThermoFisher Scientific | Cat# 12563011 |
| Gibco<sup>TM</sup> Trypan Blue Solution, 0.4% | ThermoFisher Scientific | Cat# 15-250-061 |
| Collagenase, Type I, powder | ThermoFisher Scientific | Cat # 17101015 |
| Antigen retrieval buffers used for staining:<br>citrate buffer, pH 6.0 (Sodium Citrate Dihydrate)<br>Tris-EDTA buffer, pH 9.0<br>EDTA (powder)<br>Tris base | Fisher Chemical<br>Fisher BioReagents™<br>Fisher Chemical | Cat # S279-3<br>Cat# BP118-500<br>Cat# BP154-1 |
| DAB (3,3'-Diaminobenzidine) chromogen staining | BioGenex | Cat# HK542-XAKE |
| **Critical commercial assays** |  |  |
| CXCL1/KC DuoSet ELISA kit | R & D Systems | Catalog # DY453 |
| Human IL-8/CXCL8 DuoSet ELISA | R & D Systems | Catalog # DY208 |
| Deposited data |  |  |
| https://www.ncbi.nlm.nih.gov/geo/query/acc.cgi?&acc=GSE223918. |  |  |
| **Experimental models: Cell lines** |  |  |
| 3D organoids derived from healthy human colon | This paper | NA |
| 3D organoids derived from the colon of CPC-APC APC<sup>Min/+</sup> mice. | This paper | NA |
| 2D enteroid derived monolayer (EDM) prepared from the colon of healthy human | This paper | NA |
| 2D enteroid derived monolayer (EDM) prepared from the colon of CPC-APC APC<sup>Min/+</sup> mice. | This paper | NA |
| 3D organoids derived from the colon of C567BL/6 mice. | This paper | NA |
| 2D enteroid derived monolayer (EDM) prepared from the colon of C567BL/6 mice. | This paper | NA |
| **Oligonucleotides** |  |  |
| Not applicable | Not applicable |  |
| **Recombinant DNA** |  |  |
| Not applicable | Not applicable |  |
| **Software and algorithms** |  |  |
| ImageJ | Schneider et al., 2012[105] | https://imagej.nih.gov/ij/ |
| **Other** |  |  |
| Sequence data, analyses, and resources related to the RNA sequencing of the human colon samples from healthy, non-involved region and polyp regions | This paper |  |

## Detailed Material and Methodology

### Data collection and annotation

Publicly available microarray and RNASeq databases were downloaded from the National Center for Biotechnology Information (NCBI) Gene Expression Omnibus (GEO) website[106–108]. Gene expression summarization was performed by normalizing Affymetrix platforms by RMA (Robust Multichip Average)[109, 110] and RNAseq platforms by computing TPM (Transcripts Per Millions)[111] values whenever normalized data were not available in GEO. We used log2(TPM) if TPM > 1 and (TPM—1) if TPM < 1 as the final gene expression value for analyses. We also used log2(TPM + 1) in some datasets. We also used publicly available datasets that were normalized using RPKM[112], FPKM[113, 114], TPM[115, 116], and CPM[117, 118].

#### Adenoma datasets used for network analysis

Previously published normal colon and adenoma datasets from GEO were used to perform network analysis (**Supplementary Table 1**). Samples of normal colon (N) and adenoma (A) were used for this analysis. Three validation datasets are used to test the gene signature: GSE77953 (13 N, 17 A), GSE117606/GSE117607 (65 N, 204 A) and phs001384.v1.p1 (20 N, 41 A). See **Supplementary Table 1** for all datasets analyzed in this work.

### Computational approaches

#### StepMiner analysis

StepMiner is a computational tool that identifies step-wise transitions in a time-series data[119]. StepMiner performs an adaptive regression scheme to identify the best possible step up or down based on sum-of-square errors. The steps are placed between time points at the sharpest change between low expression and high expression levels, which gives insight into the timing of the gene expression-switching event. To fit a step function, the algorithm evaluates all possible step positions, and for each position, it computes the average of the values on both sides of the step for the constant segments. An adaptive regression scheme is used that chooses the step positions that minimize the square error with the fitted data. Finally, a regression test statistic is computed as follows:

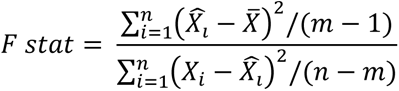

Where *X_i_* for *i* = 1 to *n* are the values, 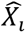 for *i* = 1 to *n* are fitted values. m is the degrees of freedom used for the adaptive regression analysis. 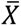 is the average of all the values: 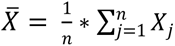. For a step position at k, the fitted values 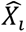 are computed by using 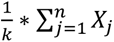 for *i* = 1 to *k* and 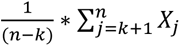 for *i* = *k* + 1 to *n*.

#### Boolean analysis

Boolean logic is a simple mathematical relationship of two values, i.e., high/low, 1/0, or positive/negative. The Boolean analysis of gene expression data requires the conversion of expression levels into two possible values. The StepMiner algorithm is reused to perform Boolean analysis of gene expression data[120]. The Boolean analysis is a statistical approach which creates binary logical inferences that explain the relationships between phenomena. Boolean analysis is performed to determine the relationship between the expression levels of pairs of genes. The StepMiner algorithm is applied to gene expression levels to convert them into Boolean values (high and low). In this algorithm, first the expression values are sorted from low to high and a rising step function is fitted to the series to identify the threshold. Middle of the step is used as the StepMiner threshold. This threshold is used to convert gene expression values into Boolean values. A noise margin of 2-fold change is applied around the threshold to determine intermediate values, and these values are ignored during Boolean analysis. In a scatter plot, there are four possible quadrants based on Boolean values: (low, low), (low, high), (high, low), (high, high). A Boolean implication relationship is observed if any one of the four possible quadrants or two diagonally opposite quadrants are sparsely populated. Based on this rule, there are six kinds of Boolean implication relationships. Two of them are symmetric: equivalent (corresponding to the positively correlated genes), opposite (corresponding to the highly negatively correlated genes). Four of the Boolean relationships are asymmetric, and each corresponds to one sparse quadrant: (low => low), (high => low), (low => high), (high => high). BooleanNet statistics is used to assess the sparsity of a quadrant and the significance of the Boolean implication relationships[45, 120]. Given a pair of genes A and B, four quadrants are identified by using the StepMiner thresholds on A and B by ignoring the Intermediate values defined by the noise margin of 2-fold change (+/- 0.5 around StepMiner threshold). Number of samples in each quadrant are defined as a_00_, a_01_, a_10_, and a_11,_ which is different from X in the previous equation of F stat. Total number of samples where gene expression values for A and B are low is computed using the following equations.

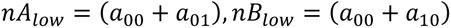

Total number of samples considered is computed using the following equation.

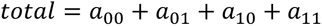

Expected number of samples in each quadrant is computed by assuming independence between A and B. For example, expected number of samples in the bottom left quadrant 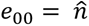 is computed as probability of A low ((*a*_00_ + *a*_01_)/*total*) multiplied by probability of B low ((*a*_00_ + *a*_10_)/*total*) multiplied by total number of samples. Following equation is used to compute the expected number of samples.

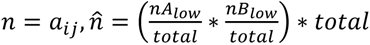

To check whether a quadrant is sparse, a statistical test for (e_00_ > a_00_) or 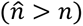 is performed by computing S_00_ and p_00_ using following equations. A quadrant is considered sparse if S_00_ is high 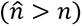 and p_00_ is small.

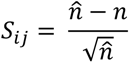

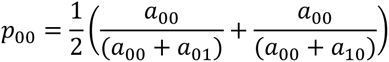

A suitable threshold is chosen for S_00_ > sThr and p_00_ < pThr to check sparse quadrant. A Boolean implication relationship is identified when a sparse quadrant is discovered using following equation.

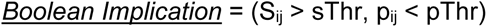

A relationship is called Boolean equivalent if top-left and bottom-right quadrants are sparse.

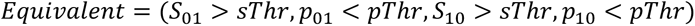

Boolean opposite relationships have sparse top-right (a_11_) and bottom-left (a_00_) quadrants.

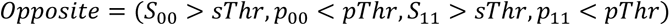

Boolean equivalent and opposite are symmetric relationships because the relationship from A to B is the same as from B to A. Asymmetric relationship forms when there is only one quadrant sparse (A low => B low: top-left; A low => B high: bottom-left; A high=> B high: bottom-right; A high => B low: top-right). These relationships are asymmetric because the relationship from A to B is different from B to A. For example, A low => B low and B low => A low are two different relationships.

A low => B high is discovered if the bottom-left (a_00_) quadrant is sparse and this relationship satisfies following conditions.

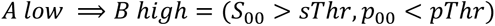

Similarly, A low => B low is identified if the top-left (a_01_) quadrant is sparse.

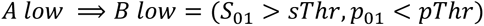

A high => B high Boolean implication is established if the bottom-right (a_10_) quadrant is sparse as described below.

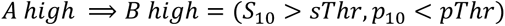

Boolean implication A high => B low is found if the top-right (a_11_) quadrant is sparse using following equation.

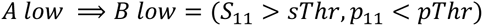

For each quadrant, a statistic S_ij_ and an error rate p_ij_ is computed. S_ij_ > sThr and p_ij_ < pThr are the thresholds used on the BooleanNet statistics to identify Boolean implication relationships.

#### Boolean network explorer (BoNE)

Boolean network explorer (*BoNE*) provides an integrated platform for the construction, visualization and querying of a network of progressive changes underlying a disease or a biological process in three steps (**Fig. S1A**): First, the expression levels of all genes in these datasets were converted to binary values (high or low) using the StepMiner algorithm. Second, gene expression relationships between pairs of genes were classified into one-of-six possible Boolean Implication Relationships (BIRs), two symmetric and four asymmetric, and expressed as Boolean implication statements. This offers a distinct advantage from conventional computational methods (Bayesian, Differential, etc.) that rely exclusively on symmetric linear relationships in networks. The other advantage of using BIRs is that they are robust to the noise of sample heterogeneity (i.e., healthy, diseased, genotypic, phenotypic, ethnic, interventions, disease severity) and every sample follows the same mathematical equation, and hence is likely to be reproducible in independent validation datasets. Third, genes with similar expression architectures, determined by sharing at least half of the equivalences among gene pairs, were grouped into clusters and organized into a network by determining the overwhelming Boolean relationships observed between any two clusters. In the resultant Boolean implication network, clusters of genes are the nodes, and the BIR between the clusters are the directed edges; *BoNE* enables their discovery in an unsupervised way while remaining agnostic to the sample type.

#### Statistical analyses

Gene signature is used to classify sample categories and the performance of the multi-class classification is measured by ROC-AUC (Receiver Operating Characteristics Area Under The Curve) values. A color-coded bar plot is combined with a density or violin+swarm plot to visualize the gene signature-based classification. All statistical tests were performed using R version 3.2.3 (2015-12-10). Standard t-tests were performed using python scipy.stats.ttest_ind package (version 0.19.0) with Welch’s Two Sample t-test (unpaired, unequal variance (equal_var=False), and unequal sample size) parameters. Multiple hypothesis corrections were performed by adjusting p values with statsmodels.stats.multitest.multipletests (fdr_bh: Benjamini/Hochberg principles). The results were independently validated with R statistical software (R version 3.6.1; 2019-07-05). Pathway analysis of gene lists were carried out via the Reactome database and algorithm [121]. Reactome identifies signaling and metabolic molecules and organizes their relations into biological pathways and processes. Kaplan-Meier analysis is performed using lifelines python package version 0.14.6.

#### Boolean implication network construction

A Boolean implication network (BIN) is created by identifying all significant pairwise Boolean implication relationships (BIRs) on a pooled dataset that included 160 normal colon and 68 adenoma samples from 11 datasets available in the human NCBI GEO Global Database for colon (**Fig. S1A**). The Boolean implication network contains the six possible Boolean relationships between genes in the form of a directed graph with nodes as genes and edges as the Boolean relationship between the genes. The nodes in the BIN are genes and the edges correspond to BIRs. Equivalent and Opposite relationships are denoted by undirected edges and the other four types (low => low; high => low; low => high; high => high) of BIRs are denoted by having a directed edge between them. The network of equivalences seems to follow a scale-free trend; however, other asymmetric relations in the network do not follow scale-free properties. BIR is strong and robust when the sample sizes are usually more than 200. The normal colon and adenoma datasets were pooled for Boolean analysis by filtering genes that had a reasonable dynamic range of expression values. When the dynamic range of expression values was small, it was difficult to distinguish if the values were all low or all high or if there were some high and some low values. Thus, it was determined that it would be best to ignore them during the Boolean analysis. The filtering step was performed by analyzing the fraction of high and low values identified by the StepMiner algorithm[119]. Any probe set or genes that contained less than 5% of high or low values were dropped from the analysis.

#### Clusters Boolean implication network

Clustering was performed in the Boolean implication network to dramatically reduce the complexity of the network (**Fig. S1C**). A clustered Boolean implication network (CBIN) was created by clustering nodes in the original BIN by following the equivalent BIRs. One approach is to build connected components in an undirected graph of Boolean equivalences. However, because of noise the connected components become internally inconsistent e.g. two genes opposite to each other becomes part of the same connected component. To avoid such a situation, we need to break the component by removing the weak links. To identify the weakest links, we first computed a minimum spanning tree for the graph and computed Jaccard similarity coefficient for every edge in this tree. Ideally, if two members are part of the same cluster, they should share as many connections as possible. If they share less than half of their total individual connections (Jaccard similarity coefficient less than 0.5) the edges are dropped from further analysis. Thus, many weak equivalences were dropped using the above algorithm, leaving the clusters internally consistent. We removed all edges that have Jaccard similarity coefficient of less than 0.5 and built the connected components with the rest. The connected components were used to cluster the BIN which is converted to the nodes of the CBIN. Increasing the Jaccard similarity cut-off will result in more compact and correlated clusters in CBIN. The distribution of cluster sizes was plotted in a log-log scale to observe the characteristics of the Boolean network (**Fig. S1D**). To ensure that the cluster sizes exhibit scale-free properties, the Jaccard similarity cut-off is modified such that they are evenly distributed along a straight line on a log-log plot (**Fig. S1D**). A new graph was built that connected the individual clusters to each other using Boolean relationships. Genes in each cluster is ranked based on the number of equivalences within the cluster. Link between two clusters (A, B) was established by using the top representative node from A that was connected to most of the members of A and, sampling 6 nodes from cluster B and identifying the overwhelming majority of BIRs (**Fig. S1C**) between the nodes from each cluster. The 6 nodes include the top representative gene (first rank), the gene next to top (second rank), middle (floor(n/2)^th^ rank where n is the cluster size), gene next to middle (floor(n/2) – 1 rank), middle from top half (floor(n/4)^th^ ranked gene), and middle from the top 1/4^th^ (floor(n/8)^th^ ranked gene) representative nodes from cluster B if size of the cluster is greater than 10. If the size of the cluster is between 2 to 10, top two and middle one are picked to test the relationship with cluster A. If the size of the cluster is 1, then it is used to test the relationship with cluster A. Testing multiple nodes provides the most common type of relationships found between clusters A and B. We suggest referring to the codebase released for additional details.

A CBIN was created using the pooled normal colon and adenoma dataset. Each cluster was associated with normal colon or adenoma samples based on where these gene clusters were highly expressed – either on the healthy or disease side. The edges between the clusters represented the Boolean relationships that are color-coded as follows: orange for low => high, dark blue for low => low, green for high => high, red for high => low, light blue for equivalent and black for opposite.

#### Boolean paths

The asymmetric BIRs provide a unique dimension to the network that is fundamentally different from any other gene expression network in the literature. Traversing a set of nodes in a directed graph of the Boolean network constitutes a Boolean path that can be interpreted as follows. A simple Boolean path involves two nodes and the directed edge between them. This simple Boolean path can be interpreted as shown in the supplementary figure (**Fig. S1E**). For the nodes X and Y with X low => Y low only quadrant #1 is sparse; the other quadrants #0, #2, and #3 are filled with samples (**Fig. S1E**). Assuming monotonicity in X and Y, the quadrants can be ordered in two possible ways: 0-2-3 and 3-2-0. The path corresponds to 0-2-3 begins with X low and Y low. This is interpreted as X turns on first and then Y turns on along a hypothetical biological path defined by the sample order. Similarly, Y turns off first and then X turns off on path 3-2-0. A complex path in the Boolean network involves more than one Boolean implication relationship (**Fig. S1F**). Three Boolean implication relationships can be used to group samples into five bins and the bins can be ordered in two possible ways (**Fig. S1F**, forward, reverse). Another example of a path is illustrated in the supplementary figure (**Fig. S1G**).

#### Discovery of paths in clustered Boolean implication network

We focus on paths that are transitive (such as **Fig. S1F-G**) because they represent a simple change in gene regulation, i.e., going from low-to-high or high-to-low once along a path (See *Boolean paths* above). By contrast, complex change refers to changes of gene regulation multiple times along a path, such as a gene going from high-to-low and then back to high. The discovery of paths starts with a node that represents the biggest cluster in the CBIN. Since a path of high=>high, high=>low, and low=>low can be used to order samples, as shown in **Fig. S1F**, we try to identify paths of this type that intersect the big clusters (top 5, based on size) in the network. To maintain the transitivity this path can be expanded as the chain of high => high, followed by high => low, followed by another chain of low => low. We would like to keep one high => low in a path because that will cover genes that are both up- and down-regulated. Since the path A high => B high can also be written as B low => A low, the chain of high => high can be reduced to the chain of low => low in the reverse direction. Therefore, we must focus only on the high => low and chain of low => low. We developed a simple, intuitive algorithm that traverses the nodes of the CBIN, starting with the biggest cluster and greedily chooses the next big cluster connected to the nodes visited in sequence. The emphasis on cluster sizes comes from the fundamental assumption that size determines importance and relevance. Therefore, we start from a big cluster (A1 from the top 5) and identify other clusters that form a chain of low => low. Further, we identify other clusters that are either opposite to A1 or they have high=>low relationship with A1, and the biggest cluster (A2) among these clusters was chosen. In addition, a chain of low=> low relationship from A2 is identified. In each subsequent step, again the biggest cluster among the different choices was greedily chosen. Finally, equivalence relationship from each cluster is used to gather more genes in each cluster and the whole path is clustered based on equivalence relationships. Depth-first traversal (DFS) was used to follow the path of low => low where bigger clusters are visited first. The search was performed until a cluster was reached for which there is no low => low relationships. For example, starting with cluster S, the search will return S low => A1 low, A1 low => A2 low, and A2 low => A3 low if A3 doesn’t have any low => low relationships. Similarly, a new starting point is considered S2 such that S2 is the biggest cluster X that has either S high => X low or S Opposite X. From cluster S2 another DFS was performed to retrieve the longest possible path of low => low. The search may return S2 low => B1 low, B1 low => B2 low if B2 doesn’t have any low => low relationships. In summary, the most prominent Boolean path was discovered by starting with the largest cluster and then exploring edges that connected to the next largest cluster in a greedy manner. This process was repeated to explore paths that connect the big clusters in the network.

#### Scoring Boolean path for sample order

A composite score was computed for a specified Boolean path that can be used to order the sample which was consistent with the logical order. To compute the score, first the genes present in each cluster were normalized and averaged. Gene expression values were normalized according to a modified Z-score approach centered around StepMiner threshold (formula = (expr -(SThr+0.5))/3*stddev). Weighted linear combination of the averages from the clusters of a Boolean path was used to create a score for each sample. The weights along the path either monotonically increased or decreased to make the sample order consistent with the logical order based on BIR. The samples were ordered based on the final weighted (RNA-seq C#1-2-3-4-5: -5 for C#1, -0.3 for C#2, 0.1 for C#3, 2.9 for C#4 and -4 for C#5; qRT-PCR: -1 for PRKAA2 and CHGA, and +1 for IL-8, LGR5, CEMIP and CLDN2; RNA-seq C#4: +1 for C#4) and linearly combined score. The direction of the path is derived from the connection from a normal colon cluster to an adenoma cluster. The sample order is visualized by a color-coded bar plot and a violin+swarm plot. A noise margin is computed for this composite score which follows the same linearly weighted combined score on 2-fold change (+/- 0.5 around StepMiner threshold).

#### Summary of genes in the clusters

Reactome pathway analysis of each cluster along the top continuum paths was performed to identify the enriched pathways[121]. The pathway description was used to summarize at a high-level what kind of biological processes are enriched in a cluster. List of genes and the pathways enriched in them are provided in Supplementary Table 2.

##### Cross-species gene name conversion

Orthologous human and mouse genes were identified using ensemble GRCh38.p13-100 gene annotations. Human to mouse gene name conversion and vice-versa used this database.

#### Charting the disease path from the normal colon to adenomas

*BoNE* uses Boolean implication network on macrophage dataset to build a signature for normal colon to adenoma disease progression. Selected clusters by size connected by high => high (green arrow), high => low (red arrows) and low => low (blue arrows) Boolean implication relationships. Reactome analysis of each clusters shows the biological processes the genes are involved in. A path is selected in the network that is used to test normal colon vs adenoma classification.

#### Boolean analysis to chart the disease path from the normal colon to adenomas

We implement supervised learning in which we use labelled training data of normal and adenoma states to train a model that can recognize a continuum of disease states. Briefly, to identify gene regulatory changes during normal colon to adenoma, we employed the MiDReG (Mining Developmentally Regulated Genes) algorithm, which utilizes statistical learning techniques.[45, 46] By applying statistical model checking to Boolean invariant rules within a static cross-sectional dataset, MiDReG infers the underlying temporal events. It identifies temporal logical changes in gene regulation by exploring transitive Boolean paths. We applied the MiDReG algorithm to analyze large and diverse normal colon and adenoma datasets (Pooled GEO; 160 normal colon, 68 adenoma), discovering Boolean invariant rules and constructing a clustered Boolean Implication Network. The model was trained by using labels from GSE76987 with normal colon (n = 41), adenoma (n = 41). The algorithm takes the adenoma network (selected graph) and this labeled dataset GSE76987 as inputs and identifies the best model to recognize the labels. The algorithm searches for transitive Boolean paths that contain three nodes with one high => low relationship. The high => low relationships cover both up/down-regulated genes and the additional Boolean path of high => high or low => low provides features to improve predictions. An unbiased search for these patterns results in 12 different Boolean paths [6,10,11], [10,11,12], [11,12,2], [1,2,3], [2,3,4], [6,7,8], [14,9,5], [12,2,4], [7,8,9], [1,2,4], [12,13,4], [14,15,4]. The nodes on the high => high side were assigned negative weights (−2, −1, etc.) and the nodes on the low => low side assigned positive weights (1, 2 etc.) to compute an optimal composite score. Two additional paths [1,2,3,4,5] and [1,2,3,4] were considered with weights from linear regression. Multivariate analysis was used to identify the best model. The performance of the Boolean path 1-2-3-4-5 was better than that of all the other paths.

#### Generation of heat maps using gene clusters identified by Boolean analysis

To generate a heatmap, a Boolean path was first constructed by following the largest clusters in the Boolean Network. Genes along this path were selected to generate a heatmap that shows the gene expression values in different samples. Gene expression values were normalized according to a modified Z-score approach centered around StepMiner threshold (formula = (expr – (SThr+0.5))/3*stddev). The samples are ordered according to an average of the normalized gene expression values in the largest cluster along the Boolean path. The heatmap uses red colors for the high values, white colors for the intermediate values and blue colors for low values. The selected genes are displayed on the left of the map.

### Human subjects

To assess the invariant gene signature and develop CRC patient-derived organoids, hereditary predisposed polyposis patients were enrolled at pediatric department at Rady Children Hospital and collection of colon biopsies from non-involved (NI) and polyp region for each patient was done following a research protocol compliant with the Human Research Protection Program (HRPP) and approved by the Institutional Review Board (Project ID#130794). Part of the patient colon biopsies was sent to the pathology lab to assess the presence/ absence of adenoma and degree CRC. In addition, the genetic mutation was assessed in each samples. Other data was gathered from each patient, such as age, gender, and previous history of the disease, following the rules of HIPAA. Each patient signed informed consent as an approval of the collection of colonic tissue biopsies for research purposes to generate 3D organoids. The colonic sample was processed for isolation and biobanking of organoids at the UC San Diego HUMANOID Center of Research Excellence (CoRE) (IRB: Project ID # 190105). The part of the colon samples was also lysed directly and RNA from the tissues was used to assess the gene expression signatures. The study design and the use of human study participants was conducted in accordance with the criteria set by the Declaration of Helsinki.

For immunohistochemical (IHC) analysis of human tissue specimens, archived formaldehyde-fixed paraffin-embedded (FFPE) human colonic biopsies from healthy controls or patients with adenomas and/or carcinomas were obtained from the Gastroenterology Division, VA San Diego (IRB# 1132632).

### Animal studies

Colon tissues were collected from NI and polyp regions of WT C57BL/6 and *CPC-APC*^Min^ mice. Part of the colon samples were lysed directly to isolate RNA for assessment of invariant gene signatures. The other part of colon samples was used to isolate intestinal crypts for enteroid isolation. The previously validated *CPC-APC* (*APC*^Min^*/+CDX2-Cre*) and control littermate mice were a kind gift from Michael Karin (UCSD); these mice are known to develop overt polyposis in the colon around the age of∼3-4 months [69, 122]. Animals were bred, housed and euthanized according to all University of California San Diego Institutional Animal Care and Use Committee (IACUC) policies under the animal protocol number S18086.

### Bacterial culture

*Fusobacterium nucleatum* (ATCC-25586) was cultured in the chopped cooked meat media in an anaerobic chamber, including an anaerobic gas kit generating system (MGC, AnaeroPACK System, Japan).

### Development of 3D colon organoids from mouse and human colons

Stem cells were isolated from the colonic crypts of mouse and human tissue specimens by digesting with Collagenase type I [2 mg/ml; Life Technologies Corporation, NY) and cultured in stem-cell enriched conditioned media (CM) containing WNT 3a, R-spondin and Noggin as described previously [104, 123–125].

### Preparation of enteroid-derived monolayers (EDMs) and infection with microbes

Enteroid-derived monolayers (EDMs) were prepared from 3D colon spheroids isolated from non-involved regions of CPC-APC^Min/+^ mice as previously described [104, 126]. Briefly, the organoids were digested with trypsin and the isolated filtered cells were resuspended in 5% CM and plated with matrigel in 0.4 μm polyester transwell membrane (Corning, Cat #3470) at the density of 2x10^5^ cell/ well. EDMs were differentiated for 2 days, and then were challenged with live microbes *Fusobacterium nucleatum (Fn)* at a multiplicity of infection 1:100 as described before [126]. The supernatant was collected from the basolateral and apical part of the transwell for cytokine analysis, and the cells were lysed for RNA extraction, followed by an analysis of target gene expression by qPCR.

### Immunohistochemistry for CEMIP in organoids

Thick sections of 4 µm formalin-fixed, paraffin-embedded (FFPE) tissues were cut and placed on glass slides coated with poly-L-lysine, followed by deparaffinization and hydration. We used citrate buffer (pH-6) and a pressure cooker to perform the heat-induced epitope step. Tissue/ organoid sections were incubated with 3% hydrogen peroxidase for 10-15 min to block endogenous peroxidase activity, followed by incubation with primary antibodies for 30-90 min in a humidified chamber at room temperature. Rabbit polyclonal anti-KIAA1199 antibody [1:50] was used for immunostaining of the organoids. We used a labeled streptavidin-biotin using 3,3′-diaminobenzidine as a chromogen and hematoxylin for visualization.

### Immunofluorescehnce of FFPE-embedded patient-derived organoids for Claudin2

FFPE organoids were placed on glass slides, followed by deparaffinization and hydration. We did the antigen retrieval step as described in IHC, then the slides were incubated with a blocking buffer at room temperature for 1 hr. Rabbit polyclonal anti-Claudin 2 antibody (1:500, Abcam) was incubated with the slides at 4⁰C for overnight. DAPI (1:1000) was used for Dapi staining. Goat anti-rabbit Alexa 594 (1:500 dilution) was used as secondary antibodies for Claudin 2. We used Leica CTR4000 Confocal Microscope for image capture, Image J software for image processing, and Illustrator software (Adobe) for image assemble and presentation.

### RNA extraction and quantitative-RT-PCR

Total RNA was extracted from NI and polyp colon biopsies of genetically predisposed CRC patients using RNA MiniPrep Kit (Zymo Research, USA) according to the manufacturer’s instructions. RNA was isolated from patient-derived organoids, APC mouse organoids, 2D EDMs challenged with different microbes using the Quick-RNA MicroPrep Kit (Zymo Research, USA) according to the manufacturer’s instruction. cDNA was prepared by the cDNA mastermix (qScript™, Quantabio). Quantitative RT-PCR (RT-qPCR) was carried out using 2x SYBR Green qPCR Master Mix (Biotool™, USA), and the cycle threshold (Ct) of target genes was normalized to 18s rRNA gene and/or β-actin. The fold change in the mRNA expression was determined using the ΔΔC_t_ method. Primers used in qPCR reactions were designed using NCBI Primer Blast software and Roche Universal Probe Library Assay Design software (Table 1).

### The sequences of primers used in Quantitative PCR reaction

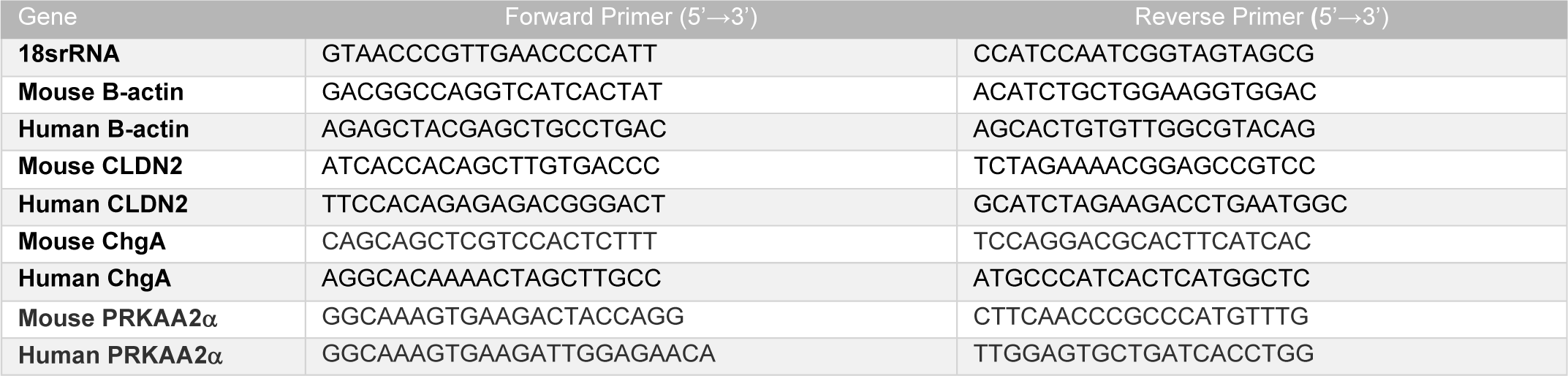
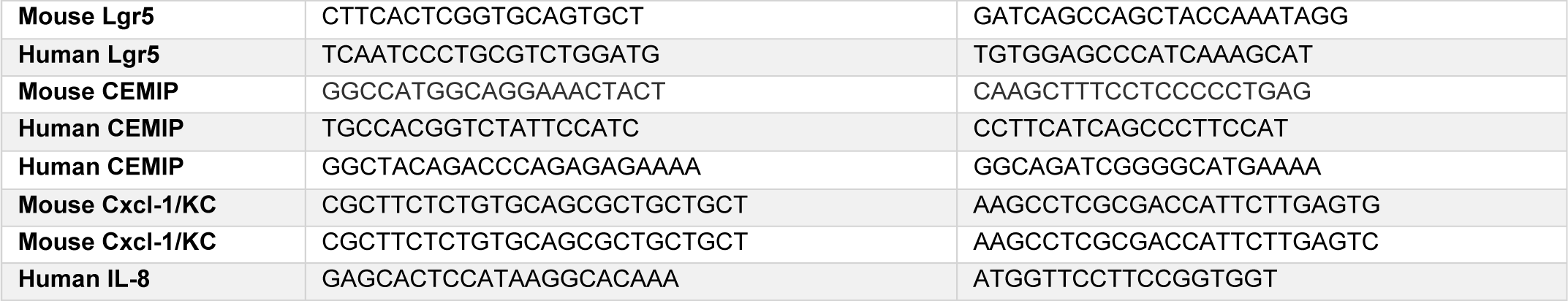

#### Quantification of mouse Cxcl-1 and human IL-8 by ELISA

Supernatants collected from organoid culture media of genetically predisposed patients and from the basolateral part of uninfected and infected monolayer were tested for IL-8 using Human IL-8 ELISA kit (Biolegend, San Diego, CA)) and Cxcl-1/KC DuoSet ELISA kit (R&D Systems, Minneapolis, MN), respectively according to the manufacturer’s instructions. IL-8 levels were compared between healthy organoids and PDO and between uninfected EDMs and infected EDMs from CPC-APC^Min/+^.

#### Statistical analyses

Data are expressed as the mean +/- S.E.M, unless stated otherwise. Statistical significance is determined using Mann-Whitney, Student’s t-test and one-way ANOVA test. *p* value is ≤ 0.05 considers significant. The graphs were generated using GraphPad version 8. Statistical tests used in each data is indicated in each figure. Statistical tests were performed using R version 3.2.3 (2015-12-10) and sklearn python packages. Heatmaps are generated using python matplotlib packages.

## Abbreviations

EDMs: Enteroid-derived monolayers
Fn: *Fusobacterium nucleatum*
PDOs: patient-derived organoids
CRCs: colorectal cancers
FAP: familial adenomatous polyposis
PJS: Peutz-Jeghers Syndrome
JPS: Juvenile Polyposis Syndrome

