## Supplement for "Molecular Signatures for Microbe-Associated Colorectal Cancers"

### Supplements

### Supplementary Tables

**Supplementary Table 1- List of datasets used for analysis**

**Supplementary Table 2- Genes identified in each cluster**

**Supplementary Table 3-Host genetics and the associated risk**

**Supplementary Table 4: Characteristics of patients used in this study**

| **Supplementary Table 1: Inventory of all publicly available gene expression datasets analyzed in this work.** | | | | |
|---|---|---|---|---|
| #Samples | Sample Type | Desc | Source | Species |
| (8, 15) | (Normal, Adenoma) | Training: Network | GSE4183 | Hs |
| (17, 0) | (Normal, Adenoma) | Training: Network | GSE11831 | Hs |
| (10, 0) | (Normal, Adenoma) | Training: Network | GSE4107 | Hs |
| (32, 32) | (Normal, Adenoma) | Training: Network | GSE8671 | Hs |
| (16, 0) | (Normal, Adenoma) | Training: Network | GSE18105 | Hs |
| (18, 0) | (Normal, Adenoma) | Training: Network | GSE9254 | Hs |
| (12, 0) | (Normal, Adenoma) | Training: Network | GSE9348 | Hs |
| (4, 0) | (Normal, Adenoma) | Training: Network | GSE13471 | Hs |
| (6, 6) | (Normal, Adenoma) | Training: Network | GSE15960 | Hs |
| (3, 5) | (Normal, Adenoma) | Training: Network | GSE10714 | Hs |
| (34, 10) | (Normal, Adenoma) | Training: Network | GSE20916 | Hs |
| (41, 41) | (Normal, Adenoma) | Training: Machine Learning | GSE76987 | Hs |
| (13, 17) | (Normal, Adenoma) | Validation | GSE77953 | Hs |
| (65, 204) | (Normal, Adenoma) | Validation | GSE117606/GSE117607 | Hs |
| (20, 41) | (Normal, Adenoma) | Validation | phs001384.v1.p1 | Hs |
| (6, 9) | (CFP, CAP) | Polyps | phs001384.v1.p1 | Hs |
| (5, 4, 11) | (Control, qUC, nUC) | UC | GSE37283 | Hs |
| (5, 5) | (Uninfected, Fn-infected) | Caco2 cells | GSE102573 | Hs |
| (9, 9) | (Normal, Fn-infected Tumor) | Colon | SRP007584 | Hs |
| (30, 19) | (Normal, Adenoma) | Validation | GSE24713 | Hs |
| (54, 51) | (Normal, Adenoma) | Validation | GSE41258 | Hs |
| (7, 10) | (Normal, Adenoma) | Validation | GSE74843 | Hs |
| (9, 11) | (Normal, Adenoma) | Validation | GSE79462 | Hs |
| (12, 24) | (Normal, Adenoma) | Validation | GSE111156 | Hs |
| (27, 32) | (Normal, Adenoma) | Validation | GSE94919 | Hs |
| (6, 6) | (Normal, Adenoma) | Validation; APC(Min/+) | GSE784 | Mm |
| (6, 6) | (Normal, Adenoma) | Validation; APC(Min/+) | GSE422 | Mm |
| (12, 10) | (Normal, Adenoma) | Validation; GEMM | GSE50794 | Mm |
| (2, 3) | (Uninfected, Wnt-independent) | Validation; colorectal cancer organoids (pool) | GSE140929 | Mm |
| (4, 6) | (Uninfected, Wnt-independent) | Validation; colorectal cancer organoids (pool+clone) | GSE140929 | Mm |
| (2, 3, 2, 3) | (Un pool, WI pool, Un clone, WI clone) | Validation; colorectal cancer organoids (pool and clone) | GSE140929 | Mm |

| | | | | |
|---|---|---|---|---|
| (3, 3, 3) | (Control, *L. casei, B. breve*) | Caco2 cells (GPL571) | GSE7259/GSE37369 | Hs |
| (6, 6, 6) | (Control, K12, O157) | Caco-2 Treatment (GPL570) | GSE50040 | Hs |
| (2, 2, 2, 2) | (Control, M90T, OspF, OspF Comp) | Caco-2 Treatments (GPL96) | GSE6082 | Hs |

| Supplementary Table 2: Genes identified in each cluster | | | | |
|---|---|---|---|---|
| MACS Path: C1-2-3-4-5 | | | | |
| C1 weight:-5 | C2 weight:-0.3 | C3 weight:0.1 | C4 weight:2.9 | C5 weight:-4 |
| PRKAA2 | FERMT2 | MALL | RAD18 | ECM1 |
| TMEM200B | RAB34 | LOC400960 | TACSTD2 | TPD52L1 |
| LAYN | ROBO1 | ENDOD1 | C1orf59 | RPP25 |
| CD109 | ATP8B2 | CPNE8 | ASCL2 | TLR4 |
| CFL2 | PTPRM | CCDC68 | LOC730101 | APCDD1 |
| DCLK1 | SDC2 | GUCA2A | REXO2 | AGT |
| TCEAL7 | ABI3BP | VIPR1 | HOXB9 | ALDH1B1 |
| CLGN | FYB | TMCC3 | PTPN13 | BRIP1 |
| PDE1A | WIPF1 | PDXP | ARHGEF10 | TTYH3 |
| FLJ25076 | MAGEH1 | LOC646627 | TRIP6 | KIF2C |
| SOX7 | JAM2 | ACAA1 | PAFAH1B3 | FUT8 |
| NAP1L3 | ZNF304 | CHP2 | RAB15 | KLK11 |
| BEX1 | ADH1B | SULT1A2 | SLC36A4 | TNS4 |
| RASSF8 | IL10RA | ITPKA | DPP7 | CCL24 |
| SDPR | ASPA | GCNT3 | CCNB1IP1 | OXGR1 |
| PEG3 | ASPN | HSD17B2 | LOC728568 | ODAM |
| | C2orf12 | SLC26A3 | CDK4 | KLK12 |
| | RECK | LRRC19 | MMP7 | KCNN4 |
| | DLC1 | ATP2B1 | ZNF473 | MGC11082 |
| | GIMAP6 | MXD1 | ZNF703 | GRIN2D |
| | VCAM1 | PTPRR | FOXQ1 | CADPS |
| | SPG20 | GUCA2B | RCC1 | LGR6 |
| | CNRIP1 | SECTM1 | PSMG4 | ETV4 |
| | HEG1 | MGC4172 | RUVBL1 | SNTB1 |
| | CADM1 | PTPRH | OSBPL3 | TCN1 |
| | CD2 | C7orf10 | KIAA1549 | FDXR |
| | CRISPLD2 | STAP2 | LOC652993 | FAIM2 |
| | PKD2 | KIF16B | SORD | RPESP |
| | AP1S2 | IL6R | TRIB3 | GPC4 |
| | FAM126A | GDPD3 | QPCT | |
| | ABCA8 | KIAA1211 | CYP4X1 | |
| | NLRC3 | HIST1H1C | GLMN | |
| | ARHGAP25 | CCL14 | GEMIN5 | |
| | FLI1 | MEP1A | 7A5 | |
| | KLRB1 | SLC4A4 | AZGP1 | |
| | C7orf58 | RAPGEFL1 | TRIM16 | |
| | MAN1C1 | CDKN2B | LGR5 | |

| | | | |
|---|---|---|---|
| | ZEB1 | TUBAL3 | CLCC1 |
| | GNB4 | FLVCR2 | TBC1D16 |
| | STMN2 | EDN3 | PHKA1 |
| | OLFML1 | PRDX6 | B9D1 |
| | SAMSN1 | SPPL2A | ZNF259 |
| | SETBP1 | SLC17A4 | CLDN1 |
| | RASSF5 | LAMA1 | MTERFD3 |
| | EMCN | C17orf76 | RIPK2 |
| | RCAN2 | IGSF9 | MET |
| | TMEM204 | SLC25A20 | CD44 |
| | RDX | RND3 | WDR77 |
| | PCDH7 | ACSS2 | MYC |
| | MYO5A | RHOF | SLC7A5 |
| | CLIP4 | LOC400573 | CAD |
| | LDB2 | C1orf106 | LOC645166 |
| | ARMCX1 | PIGZ | SLC6A6 |
| | GMFG | GCNT2 | SLC29A1 |
| | ARHGEF6 | SULT1A1 | PDCD2L |
| | LIFR | TP53INP2 | KLHL29 |
| | GPC6 | CNNM4 | ABCC1 |
| | P2RY14 | GGT6 | CDC25B |
| | GALNAC4S-6ST | SRI | ALS2CR4 |
| | CYYR1 | OAF | RNASEH2A |
| | IL18BP | MGC13057 | CEP78 |
| | PTPLAD2 | KCNK5 | KIAA1199 |
| | GIMAP7 | SCIN | ARNTL2 |
| | HERC5 | SAMD9 | C1orf67 |
| | FRMD6 | PDLIM2 | PROX1 |
| | RCSD1 | PCK1 | XPO5 |
| | CYBRD1 | NHSL1 | CBR3 |
| | SSBP2 | TMEM45B | EPHB2 |
| | COL14A1 | ENTPD5 | SP5 |
| | OLFML3 | TMEM120A | JUB |
| | ITPR1 | TRPM6 | OTUB2 |
| | MAMDC2 | TMEM37 | FAM152B |
| | SLC2A5 | C14orf139 | C6orf125 |
| | LBH | CPM | VSNL1 |
| | RGL1 | RP11-285G1.3 | COPG2 |
| | WWTR1 | SCNN1B | ZNRF3 |

| | | | |
|---|---|---|---|
| | TMEM47 | LOC100133660 | NFE2L3 |
| | PALM2-AKAP2 | MMP28 | DACH1 |
| | GIMAP4 | SLC36A1 | TBX3 |
| | C10orf128 | CA4 | GALNT6 |
| | DOCK10 | TSPAN1 | TEAD4 |
| | DPYD | CA2 | ATP11A |
| | SCPEP1 | PLA2G10 | SLC39A10 |
| | AKT3 | PADI2 | S100A2 |
| | MITF | ABCG2 | IFITM2 |
| | CSRP2 | SLC22A18AS | IL8 |
| | PTPRC | CA1 | ICA1 |
| | MEF2C | RELL1 | CLDN2 |
| | GIMAP8 | CMBL | RNF43 |
| | LOC283666 | CA7 | RGNEF |
| | PPP1R16B | STBD1 | GTF2IRD1 |
| | ITGA4 | PRSS8 | LOC100129762 |
| | SGCE | CLDN23 | LRRC6 |
| | SYNE1 | LPAR1 | LPCAT1 |
| | KIAA1946 | CNNM2 | DGAT2 |
| | FLRT2 | ATP1B3 | KRT80 |
| | C14orf132 | SLC25A34 | TESC |
| | MAP1B | TST | PPM1H |
| | TSPYL5 | SDCBP2 | TDGF1 |
| | HCLS1 | LOC100130886 | SLC35E4 |
| | GIMAP1 | CLIC5 | REPS2 |
| | PDE7B | C10orf54 | FXN |
| | HLA-DMB | AGPAT9 | LOC652900 |
| | CXCL12 | SMPDL3A | TRAP1 |
| | LY86 | CLCN2 | RAD54B |
| | TEK | GRAMD3 | FAM92A1 |
| | MAFB | MEIS1 | TGFBI |
| | CLEC2B | CASP7 | CDH3 |
| | MSRB3 | KRT20 | AXIN2 |
| | CCDC88A | TSPAN7 | LOC254057 |
| | PMP22 | RHOU | EPHB3 |
| | RBMS1 | ITM2C | LOC100134295 |
| | FBN1 | DDX60 | FGGY |
| | AKR1B1 | PEX26 | CCDC113 |
| | FILIP1L | ALPI | FAM148A |
| | AKAP12 | TMEM171 | KIF18A |

| | | | |
|---|---|---|---|
| | TRAC | OSTbeta | HIST3H2A |
| | DSE | SPIB | SLC38A5 |
| | LAMA4 | HSD11B2 | MRE11A |
| | DCN | HIGD1A | CYB5R2 |
| | EVI2A | DHRS9 | NOB1 |
| | JAZF1 | CES2 | ASNS |
| | FZD1 | PKIB | |
| | CFH | AHCYL2 | |
| | ZEB2 | PPAP2A | |
| | RASSF2 | MS4A12 | |
| | CD48 | SLC22A5 | |
| | FAM129A | SGK2 | |
| | CLIC2 | RUNDC3B | |
| | EVI2B | SEMA6D | |
| | MAF | | |
| | RHOJ | | |
| | RFTN1 | | |
| | LMO2 | | |
| | NKX2-3 | | |
| | ARHGAP15 | | |
| | APBB1IP | | |
| | ANK2 | | |
| | QKI | | |
| | C1orf54 | | |
| | PGCP | | |
| | DOCK2 | | |
| | EFEMP1 | | |
| | ARHGAP30 | | |
| | MCC | | |
| | DARC | | |
| | SCARA5 | | |
| | LIX1L | | |
| | CD27 | | |
| | GNG2 | | |
| | RBMS3 | | |
| | SRPX | | |

| MACS Signature: C4 | Ranking of C4 genes based on T-test between Normal and Adenoma in Pooled Dataset | | | | |
|---|---|---|---|---|---|
| C4 weight:1 | ID | Name | T | P | Log2FC |
| RAD18 | 212942_s_at | KIAA1199 | -50.029244 | 9.94E-73 | -4.4893377 |
| TACSTD2 | 227475_at | FOXQ1 | -25.418741 | 1.47E-40 | -4.1834445 |
| C1orf59 | 227702_at | CYP4X1 | -14.732652 | 3.75E-23 | -3.6622361 |
| ASCL2 | 202286_s_at | TACSTD2 | -19.874737 | 9.27E-34 | -3.6028096 |
| LOC730101 | 241031_at | FAM148A | -14.748229 | 3.30E-24 | -3.2850442 |
| REXO2 | 204259_at | MMP7 | -18.439727 | 1.29E-28 | -3.2281878 |
| HOXB9 | 213880_at | LGR5 | -18.862805 | 3.39E-36 | -3.1058674 |
| PTPN13 | 222696_at | AXIN2 | -27.038259 | 3.21E-59 | -3.0413459 |
| ARHGEF10 | 229215_at | ASCL2 | -20.622931 | 3.16E-35 | -2.8468888 |
| TRIP6 | 223509_at | CLDN2 | -19.912716 | 2.75E-31 | -2.8236625 |
| PAFAH1B3 | 222549_at | CLDN1 | -18.722349 | 1.41E-31 | -2.7054453 |
| RAB15 | 206286_s_at | TDGF1 | -20.882906 | 1.74E-42 | -2.5384493 |
| SLC36A4 | 219956_at | GALNT6 | -21.761563 | 2.69E-39 | -2.5120328 |
| DPP7 | 219682_s_at | TBX3 | -19.611264 | 2.61E-34 | -2.5002265 |
| CCNB1IP1 | 203256_at | CDH3 | -21.41009 | 8.47E-39 | -2.398607 |
| LOC728568 | 202859_x_at | IL8 | -9.8606011 | 1.96E-17 | -2.3484831 |
| CDK4 | 228915_at | DACH1 | -15.108502 | 5.82E-29 | -2.3070511 |
| MMP7 | 225806_at | JUB | -15.419841 | 2.01E-26 | -2.29674 |
| ZNF473 | 205174_s_at | QPCT | -14.530524 | 7.06E-26 | -2.2807836 |
| ZNF703 | 201506_at | TGFBI | -19.881972 | 7.44E-40 | -2.2689525 |
| FOXQ1 | 228754_at | SLC6A6 | -18.405305 | 2.11E-35 | -2.2603242 |
| RCC1 | 226360_at | ZNRF3 | -17.517534 | 3.40E-38 | -2.1681637 |
| PSMG4 | 201563_at | SORD | -20.295248 | 6.58E-51 | -2.156211 |
| RUVBL1 | 204702_s_at | NFE2L3 | -19.128137 | 5.73E-35 | -2.1147791 |

| | | | | | |
|---|---|---|---|---|---|
| OSBPL3 | 209835_x_at | CD44 | -16.248154 | 2.97E-33 | -2.0525156 |
| KIAA1549 | 209309_at | AZGP1 | -14.788241 | 1.76E-25 | -2.0492238 |
| LOC652993 | 212686_at | PPM1H | -20.806516 | 7.62E-41 | -1.9716072 |
| SORD | 201195_s_at | SLC7A5 | -16.878816 | 1.60E-31 | -1.9439268 |
| TRIB3 | 218704_at | RNF43 | -22.603712 | 1.18E-40 | -1.8921061 |
| QPCT | 202431_s_at | MYC | -14.064346 | 6.77E-28 | -1.8081099 |
| CYP4X1 | 203510_at | MET | -17.34115 | 1.14E-38 | -1.7694252 |
| GLMN | 1438_at | EPHB3 | -11.90169 | 4.98E-20 | -1.7109562 |
| GEMIN5 | 203798_s_at | VSNL1 | -12.830632 | 5.04E-21 | -1.6976883 |
| 7A5 | 228656_at | PROX1 | -10.571876 | 2.76E-17 | -1.6739625 |
| AZGP1 | 212806_at | LOC100129762 | -12.336786 | 1.61E-21 | -1.6514641 |
| TRIM16 | 218412_s_at | GTF2IRD1 | -23.557046 | 5.07E-45 | -1.6057514 |
| LGR5 | 219494_at | RAD54B | -15.39964 | 8.67E-28 | -1.5935512 |
| CLCC1 | 219911_s_at | LOC100134295 | -14.314115 | 1.71E-24 | -1.591636 |
| TBC1D16 | 232151_at | 7A5 | -17.000877 | 4.38E-31 | -1.5820053 |
| PHKA1 | 230875_s_at | ATP11A | -10.120108 | 8.55E-17 | -1.5780662 |
| B9D1 | 209627_s_at | OSBPL3 | -17.23808 | 3.58E-34 | -1.5563918 |
| ZNF259 | 201801_s_at | SLC29A1 | -12.568184 | 1.58E-21 | -1.5401055 |
| CLDN1 | 201853_s_at | CDC25B | -13.647922 | 1.14E-23 | -1.500063 |
| MTERFD3 | 204268_at | S100A2 | -10.893168 | 4.77E-19 | -1.468869 |
| RIPK2 | 225295_at | SLC39A10 | -10.620771 | 2.66E-19 | -1.4500583 |
| MET | 222890_at | CCDC113 | -13.389942 | 2.60E-23 | -1.4426685 |
| CD44 | 231849_at | KRT80 | -13.054023 | 2.50E-21 | -1.4416742 |
| WDR77 | 232370_at | LOC254057 | -13.007802 | 1.94E-21 | -1.4336496 |
| MYC | 41037_at | TEAD4 | -14.739829 | 6.42E-26 | -1.4326031 |

| | | | | | |
|---|---|---|---|---|---|
| SLC7A5 | 218872_at | TESC | -11.155278 | 7.54E-18 | -1.3825364 |
| CAD | 218145_at | TRIB3 | -11.96436 | 1.41E-19 | -1.3580462 |
| LOC645166 | 235391_at | FAM92A1 | -10.088051 | 2.67E-16 | -1.3515291 |
| SLC6A6 | 223018_at | NOB1 | -18.701444 | 3.01E-38 | -1.3425966 |
| SLC29A1 | 204341_at | TRIM16 | -11.668811 | 9.30E-20 | -1.3194549 |
| PDCD2L | 204201_s_at | PTPN13 | -11.652342 | 1.30E-18 | -1.314285 |
| KLHL29 | 230002_at | CLCC1 | -13.025254 | 5.16E-24 | -1.2905482 |
| ABCC1 | 222116_s_at | TBC1D16 | -17.678532 | 9.03E-36 | -1.2815504 |
| CDC25B | 229310_at | KLHL29 | -10.926444 | 9.49E-18 | -1.2807452 |
| ALS2CR4 | 209589_s_at | EPHB2 | -12.217729 | 2.84E-22 | -1.2784727 |
| RNASEH2A | 224467_s_at | PDCD2L | -19.434752 | 7.26E-40 | -1.270457 |
| CEP78 | 201614_s_at | RUVBL1 | -13.921267 | 3.06E-27 | -1.2513633 |
| KIAA1199 | 227425_at | REPS2 | -10.645279 | 9.95E-19 | -1.2425804 |
| ARNTL2 | 202804_at | ABCC1 | -16.991591 | 1.07E-32 | -1.2355875 |
| C1orf67 | 205395_s_at | MRE11A | -11.101854 | 1.13E-19 | -1.2336981 |
| PROX1 | 235845_at | SP5 | -18.670008 | 7.39E-35 | -1.2240719 |
| XPO5 | 226064_s_at | DGAT2 | -15.631253 | 2.89E-28 | -1.2240605 |
| CBR3 | 213248_at | LOC730101 | -12.4545 | 6.41E-22 | -1.2025325 |
| EPHB2 | 221258_s_at | KIF18A | -9.4748778 | 3.55E-15 | -1.2004011 |
| SP5 | 220658_s_at | ARNTL2 | -12.578639 | 8.95E-22 | -1.1918989 |
| JUB | 201420_s_at | WDR77 | -18.050658 | 3.62E-35 | -1.1918168 |
| OTUB2 | 217988_at | CCNB1IP1 | -12.116074 | 4.64E-24 | -1.1601354 |
| FAM152B | 201818_at | LPCAT1 | -12.390396 | 2.36E-22 | -1.1580693 |
| C6orf125 | 218194_at | REXO2 | -10.011365 | 6.13E-17 | -1.1535549 |
| VSNL1 | 222760_at | ZNF703 | -17.049527 | 4.27E-32 | -1.1484539 |

| | | | | | |
|---|---|---|---|---|---|
| COPG2 | 232994_s_at | RGNEF | -14.963685 | 1.15E-27 | -1.1475388 |
| ZNRF3 | 209129_at | TRIP6 | -10.7301 | 1.09E-17 | -1.1473144 |
| NFE2L3 | 216417_x_at | HOXB9 | -11.223996 | 3.97E-19 | -1.1380056 |
| DACH1 | 207153_s_at | GLMN | -14.720986 | 1.38E-29 | -1.1280265 |
| TBX3 | 1568623_a_at | SLC35E4 | -13.350951 | 5.17E-23 | -1.1265455 |
| GALNT6 | 206483_at | LRRC6 | -14.90807 | 3.11E-27 | -1.122303 |
| TEAD4 | 203022_at | RNASEH2A | -9.9105475 | 4.26E-16 | -1.1070781 |
| ATP11A | 202246_s_at | CDK4 | -10.703367 | 3.03E-19 | -1.0927069 |
| SLC39A10 | 59697_at | RAB15 | -11.886495 | 2.82E-21 | -1.0791183 |
| S100A2 | 225346_at | MTERFD3 | -13.138633 | 7.05E-26 | -1.0723551 |
| IFITM2 | 201315_x_at | IFITM2 | -8.6774737 | 1.12E-13 | -1.0569559 |
| IL8 | 229876_at | PHKA1 | -12.273401 | 9.54E-21 | -1.0449705 |
| ICA1 | 224200_s_at | RAD18 | -10.872165 | 6.65E-19 | -1.0418898 |
| CLDN2 | 225841_at | C1orf59 | -11.640112 | 5.51E-20 | -1.0390865 |
| RNF43 | 210534_s_at | B9D1 | -11.076798 | 4.20E-19 | -1.036791 |
| RGNEF | 205379_at | CBR3 | -12.105533 | 1.98E-21 | -1.0339169 |
| GTF2IRD1 | 205565_s_at | FXN | -11.169712 | 1.80E-19 | -1.0280467 |
| LOC100129762 | 228217_s_at | PSMG4 | -11.041245 | 1.82E-20 | -1.0151108 |
| LRRC6 | 201391_at | TRAP1 | -12.93236 | 8.26E-23 | -0.9998286 |
| LPCAT1 | 209545_s_at | RIPK2 | -13.139982 | 2.87E-24 | -0.9911643 |
| DGAT2 | 218720_x_at | LOC652900 | -11.981233 | 2.99E-21 | -0.9804084 |
| KRT80 | 223457_at | COPG2 | -10.894497 | 2.24E-18 | -0.9797153 |
| TESC | 219718_at | FGGY | -12.895598 | 1.54E-22 | -0.9794381 |
| PPM1H | 223575_at | KIAA1549 | -13.148502 | 7.30E-23 | -0.9782701 |
| TDGF1 | 205047_s_at | ASNS | -7.4831278 | 4.22E-11 | -0.9720524 |
| SLC35E4 | 221582_at | HIST3H2A | -12.753363 | 5.87E-22 | -0.957259 |

| | | | | | |
|---|---|---|---|---|---|
| REPS2 | 217998_at | LOC652993 | -12.043865 | 8.05E-21 | -0.9565272 |
| FXN | 206499_s_at | RCC1 | -11.132859 | 1.93E-19 | -0.9457039 |
| LOC652900 | 1553956_at | ALS2CR4 | -11.012251 | 4.79E-19 | -0.9408626 |
| TRAP1 | 202715_at | CAD | -12.472893 | 1.49E-22 | -0.9372061 |
| RAD54B | 234973_at | SLC38A5 | -8.8496589 | 8.17E-14 | -0.9356674 |
| FAM92A1 | 203228_at | PAFAH1B3 | -11.351648 | 1.26E-20 | -0.9287938 |
| TGFBI | 210547_x_at | ICA1 | -11.677055 | 3.37E-21 | -0.8910058 |
| CDH3 | 219369_s_at | OTUB2 | -10.894578 | 8.75E-18 | -0.8904202 |
| AXIN2 | 223057_s_at | XPO5 | -10.329456 | 3.60E-17 | -0.8761233 |
| LOC254057 | 238012_at | DPP7 | -10.067592 | 1.48E-16 | -0.8754043 |
| EPHB3 | 225712_at | GEMIN5 | -10.592461 | 8.66E-19 | -0.8596632 |
| LOC100134295 | 220230_s_at | CYB5R2 | -9.8008351 | 7.81E-16 | -0.84947 |
| FGGY | 200054_at | ZNF259 | -9.9673076 | 1.70E-16 | -0.8334939 |
| CCDC113 | 242283_at | C1orf67 | -9.9891688 | 3.98E-16 | -0.8330357 |
| FAM148A | 228774_at | CEP78 | -9.7946624 | 3.40E-16 | -0.7905884 |
| KIF18A | 234978_at | SLC36A4 | -8.8812669 | 1.63E-14 | -0.7769029 |
| HIST3H2A | 228158_at | LOC645166 | -6.2701862 | 1.45E-08 | -0.7688007 |
| SLC38A5 | 212527_at | FAM152B | -8.4841106 | 2.37E-13 | -0.7588751 |
| MRE11A | 226943_at | LOC728568 | -7.653796 | 7.95E-12 | -0.7471652 |
| CYB5R2 | 224448_s_at | C6orf125 | -7.7290764 | 2.41E-12 | -0.7386832 |
| NOB1 | 216620_s_at | ARHGEF10 | -7.4570279 | 5.56E-11 | -0.7259855 |
| ASNS | 213124_at | ZNF473 | -9.1772854 | 1.99E-14 | -0.7001635 |

| Core MACS Genes | |
|---|---|
| Weight:-1 | Weight: 1 |
| PRKAA2 | CXCL8/IL-8 |
| CHGA | LGR5 |
| | CEMIP |
| | CLDN2 |

| ID | Name | T | P | Log2FC |
|---|---|---|---|---|
| 238441_at | PRKAA2 | 12.722135 | 4.132351e-28 | 1.109298 |
| 204697_s_at | CHGA | 15.084556 | 5.13E-32 | 2.130831 |
| 202859_x_at | IL8 | -9.860601 | 1.961075e-17 | -2.348483 |
| 213880_at | LGR5 | -18.862805 | 3.39E-36 | -3.105867 |
| 212942_s_at | KIAA1199 | -50.029244 | 9.94E-73 | -4.489338 |
| 223509_at | CLDN2 | -19.912716 | 2.750723e-31 | -2.823662 |

### Supplementary Table 3-Host genetics and the associated risk

| DISEASE | GENE | INCIDENCE | RISK (AVG. AGE AT CRC DIAGNOSIS) | CLUSTER NUMBER | PREDOMINANT CANCER |
|---|---|---|---|---|---|
| **Familial adenomatous polyposis (FAP)** | *APC* | 1 in 7,000 to 1 in 22,000 live births. | Lifetime risk **100%** (39 years) | Cluster 6 | Colorectal, small bowel, gastric, etc. |
| **Attenuated FAP (AFAP)** | *APC* | Unknown. | Lifetime risk **70%** (56 years) | Cluster 6 | |
| **Lynch syndrome (HNPCC)** | *MLH1, MSH2, MSH6, and PMS2. EPCAM* | 1:370 to 1:2,000 in Western populations | Lifetime risk **60–80%** (45 years) | MLH1, MSH2 (Cluster 7) MSH6 (Cluster 8) | Multiple (including colorectal and endometrial) |
| **MUTYH-associated polyposis** | *MUTYH* | 1 per 10.000 and 40.000 newborns. | Lifetime risk **80%** for biallelic; 5-7% for monoallelic. (50 years; 26-98) | Cluster 6 | Colorectal |
| **Peutz-Jeghers syndrome (PJS)** | *STK11 (LKB1)* | 1 in 8300 to 1 in 280 000 individuals | Lifetime risk **39%** (fifth decade of life) | Cluster 6 | Multiple (including colorectal, small bowel, pancreas) |
| **Juvenile polyposis syndrome (JPS)** | *BMPR1A, SMAD4* | 1 in 100,000 to 160,000 individuals | Lifetime risk **40-50%**. (third decade of life) | SMAD 4 (Cluster 7) | Colorectal and gastric cancer |
| **Cowden syndrome** | *PTEN (TEP1)* | 1 case per 200,000 population | Lifetime risk **16%** (sixth decade) | Cluster 6 | Multiple (including colorectal) |
| **Li-Fraumeni syndrome** | *p53 (TP53)* | Unknown. | | Cluster 8 | Multiple (including colorectal) |
| **Sporadic colon cancer** | *APC* | 40 *per* 100,000 [68 (men) and 72 (women)] | Lifetime risk **5-6%** | Cluster 6 | Colon |
| | *KRAS* | | | Singleton | |
| | *NRAS* | | | Cluster 7 | |
| | *RRAS2* | | | Cluster 7 | |
| | *BRAF* | | | Singleton | |
| | *PIK3CA* | | | Cluster 7 | |
| | *P53* | | | Cluster 8 | |
| | *SMAD4* | | | Cluster 7 | |
| | *MLH1* | | | Cluster 7 | |

### Supplementary Table 4: Characteristics of patients used in this study

| Genotype | Patient information (mutation) | Gender | Age (years) |
|---|---|---|---|
| **H4** | Healthy | M | 60-64 |
| **H14** | Healthy | M | 45-49 |
| **H19** | Healthy | F | 45-49 |
| **FAP1** | Familial Adenomatous Polyposis, APC c.646C>T (p.Arg216Ter) | M | 8.5 |
| **FAP4** | Familial Adenomatous Polyposis, APC c.1370C>A (p.Ser457Ter) | M | 16 |
| **FAP5** | Familial Adenomatous Polyposis, APC c.643C>T (p.Gln215Ter) | F | 13.5 |
| **FAP6** | Familial Adenomatous Polyposis, APC c.643C>T (p.Gln215Ter) | F | 9 |
| **FAP7** | Familial Adenomatous Polyposis, MYH D12.6 1145 G>A; MYH 494 A>G | F | 14 |
| **Lynch 1** | Lynch Syndrome, MSH2, c.2228C>G (p.S743X) . | M | 12 |
| **Lynch 2** | Lynch Syndrome, MSH2, c.2228C>G (p.S743X). | F | 14 |
| **PJS1** | Peutz-Jeghers syndrome, SGS (short gut syndrome),Dumping syndrome, Chromosome 22q13 microdeletion syndrome | F | 16 |
| **PJS2** | Peutz-Jeghers syndrome, Duodenal mass, STK11, c.385 dupA, aa alteration pMet129fs | M | 13 |
| **JPS** | Juvenile polyposis syndrome, heterozygous for the EX4-5 del gross deletion in the BMPR1A gene | M | 14 |

** The above list indicates all the samples collected for this study. Considering the challenges and amount of tissues, we have restricted the use of every sample in every assay related to organoid isolation-functional assays, transcriptomics and immunostaining. The specifics of the disease types used in the experiments have been mentioned in the figure legends.**

### Supplementary Figures with Figure legends

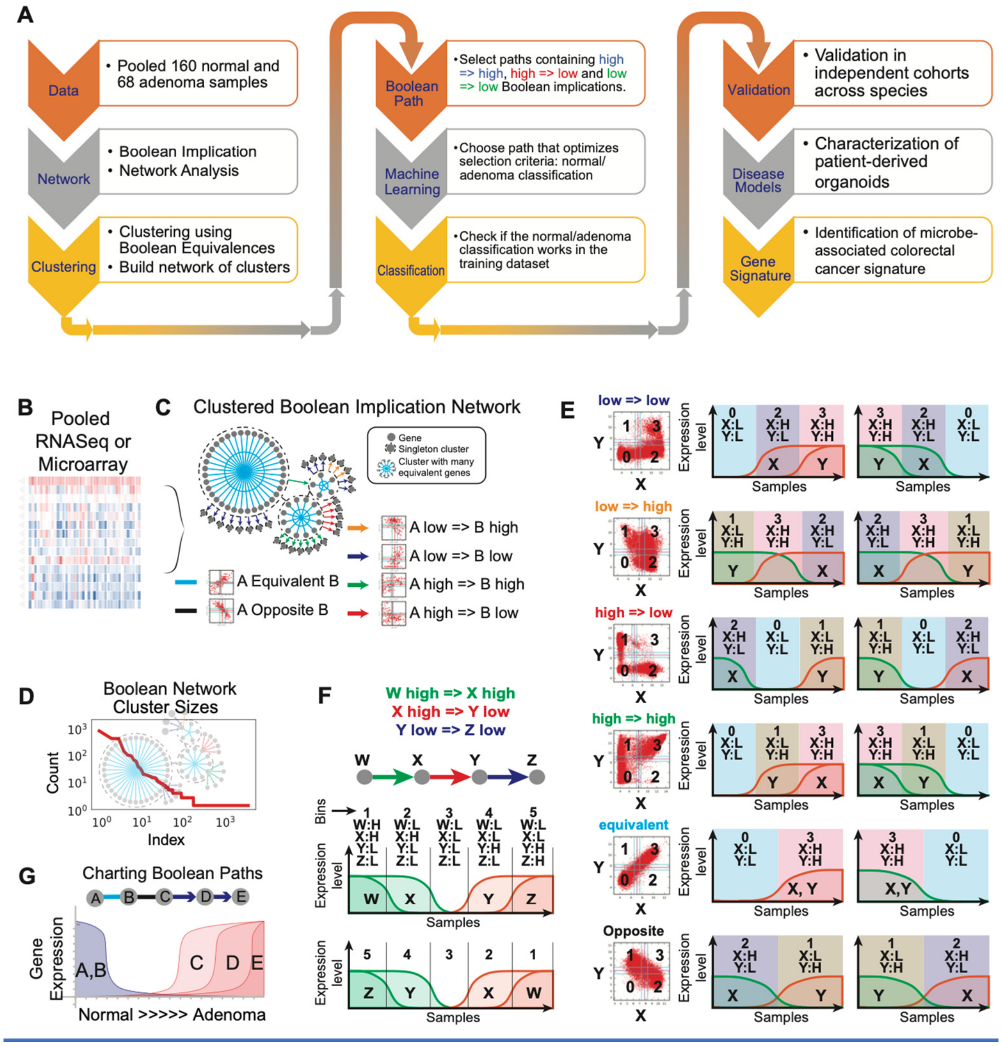

**Supplementary Figure 1: Boolean Network Explorer (BoNE): A tool for clustering and visualization of the Boolean implication network. A)** Overview of the computational steps used in BoNE. **B)** BoNE was applied to analyze normal and adenoma datasets to develop a model of polarization. A pooled dataset is used to build the Boolean implication network. **C)** BooleanNet algorithm is applied to identify Boolean implication relationships. The BoNE uses Boolean equivalent relationships to cluster genes and identify relationships between clusters. **D)** A graphical display of cluster size analysis shows a linear trend in log-log scatterplots between clusters sorted by size and the number of clusters of any size. **E)** Sample ordering based on single Boolean Implication relationships. **F)** Sample ordering based on a sequence of high => high, high => low, low => low Boolean relationships. **G)** Like panel F, four Boolean implication relationships ‘A equivalent to B’, ‘B opposite C’, ‘C low => D low’ and ‘D low => E’ low constitute a Boolean path that can be used to develop a computational model of the progression from normal to adenoma. A suitable path is selected by using machine learning that optimizes the strength of normal/adenoma classification.

#### Validation of Path #1-2-3-4-5

| Dataset | Cohort | Species | Normal (n) | Adenoma (n) | ROC AUC | |
|---|---|---|---|---|---|---|
| **GEO Pooled** | Test | Human | 160 | 68 | 1.00 | Pooled GEO |
| **GSE77953** | Validation | Human | 13 | 17 | 0.98 | GSE77953 |
| **GSE117607** | Validation | Human | 65 | 204 | 0.98 | GSE117606 / 7 |
| **phs001384.v1.p1** | Validation | Human | 65 | 204 | 1.00 | phs001384.v1.p1 |
| **GSE4183** | Validation | Human | 8 | 15 | 1.00 | GSE4183 |
| **GSE8671** | Validation | Human | 32 | 32 | 1.00 | GSE8671 |
| **GSE24713** | Validation | Human | 30 | 19 | 1.00 | GSE24713 |
| **GSE41258** | Validation | Human | 54 | 51 | 0.98 | GSE41258 |
| **GSE74843** | Validation | Human | 7 | 10 | 0.86 | GSE74843 |
| **GSE79462** | Validation | Human | 9 | 11 | 0.84 | GSE79462 |
| **GSE76987** | Test | Human | 41 | 41 | 0.85 | GSE76987 |
| **GSE111156** | Validation | Human | 12 | 24 | 0.97 | GSE111156 |
| **GSE94919** | Validation | Human | 27 | 32 | 0.86 | GSE94919 |
| **GSE102573** | Validation | Human | 5 | 5 | 1.00 | GSE102573 |
| **SRP007584** | Validation | Human | 9 | 9 | 1.00 | SRP007584 |
| **GSE784** | Validation | Mouse | 6 | 6 | 0.86 | GSE784 |
| **GSE422** | Validation | Mouse | 6 | 5 | 0.63 | GSE422 |
| **GSE50794** | Validation | Mouse | 12 | 10 | 1.00 | GSE50794 |

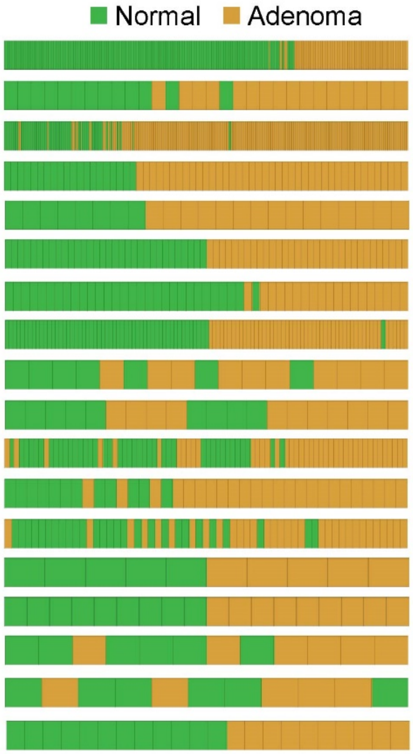

**Supplementary Figure 2:** The C#1-2-3-4-5 path separates healthy vs adenoma samples. Machine learning identified the C#1-2-3-4-5 path segregates normal vs adenoma samples in two datasets: GEO Pooled and GSE76987. The C#1-2-3-4-5 path is applied to sixteen validation datasets (13 human datasets and 3 mouse dataset) to predict normal vs adenoma samples: Human datasets are GSE77953, GSE117607, phs001384.v1.p1, GSE4183, GSE8671, GSE24713, GSE41258, GSE74843, GSE79462, GSE111156, GSE94919, GSE102573, and SRP007584. Mice datasets are GSE784, GSE422, and GSE50794. The strength of the sample separation is determined by the number of samples, ROC AUC, Accuracy, and Fisher exact p-values.

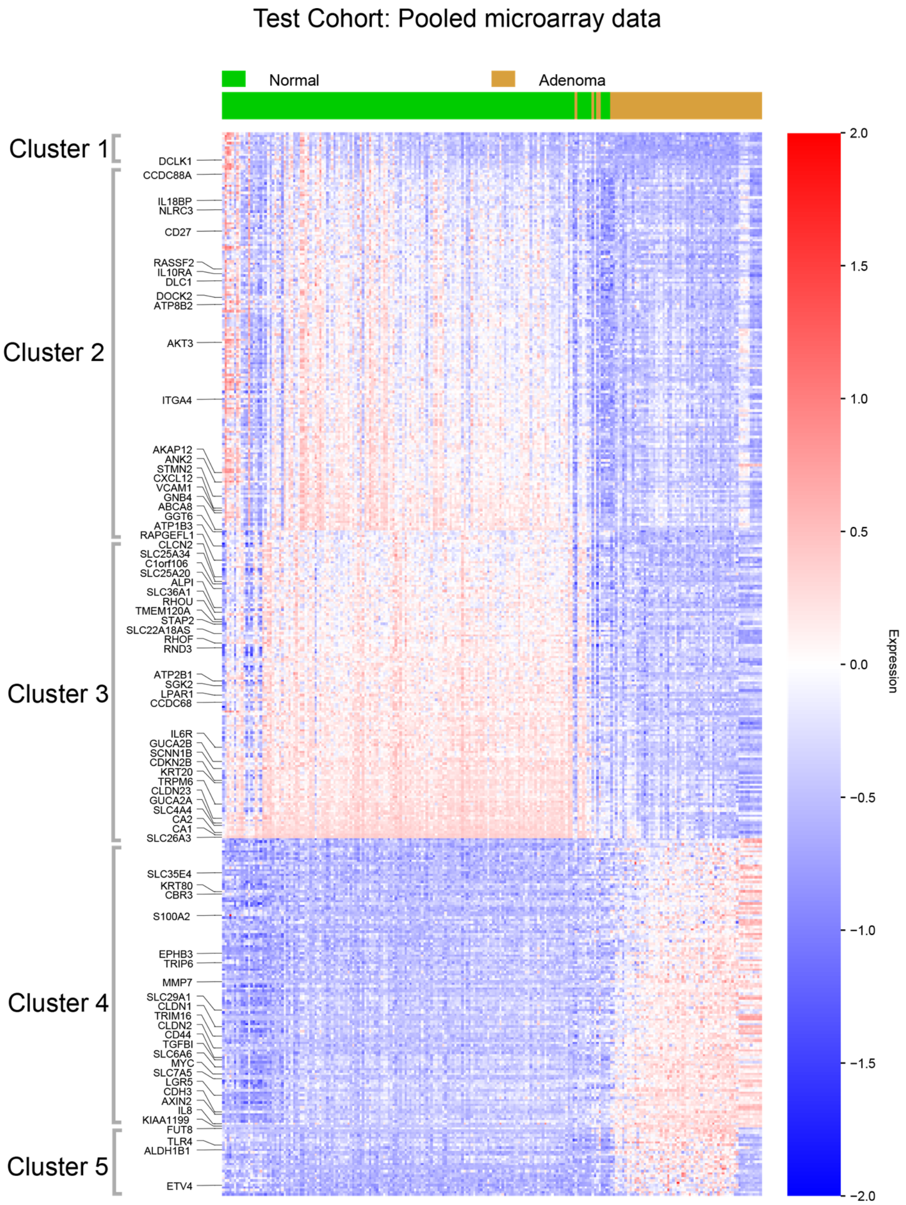

| | Precision [TP/(TP + FP)] | Recall [TP/(TP+FN)] | F1-score [2 x (precision x recall)/(precision + recall)] | Support [total samples] |
|---|---|---|---|---|
| **Normal (training)** | 0.98 | 0.99 | 0.98 | 160 |
| **Adenoma (training)** | 0.98 | 0.94 | 0.96 | 68 |

TP = True Positives; FP= False positives; FN = False negatives
**ROC AUC** (Receiver Operator Characteristic - Area under curve) = 0.998
**Classification Accuracy Score:** 0.98
**Fisher Exact p value:** 1.46 e-50

**Supplementary Figure 3:** Heatmap of genes present in the C#1-2-3-4-5 path in the test cohort. Key genes in the clusters are presented on the left of the heatmap. The expression level ranged from -1 to +1; the expression below 0 indicated low expression and represented with blue color and the expression above 0 indicated high expression and represented with red color. Sample ranking using the C#1-2-3-4-5 path is shown in the bar plot above the heatmap.

Validation Cohort #1: dbGaP Study Accession: phs001384.v1.p1

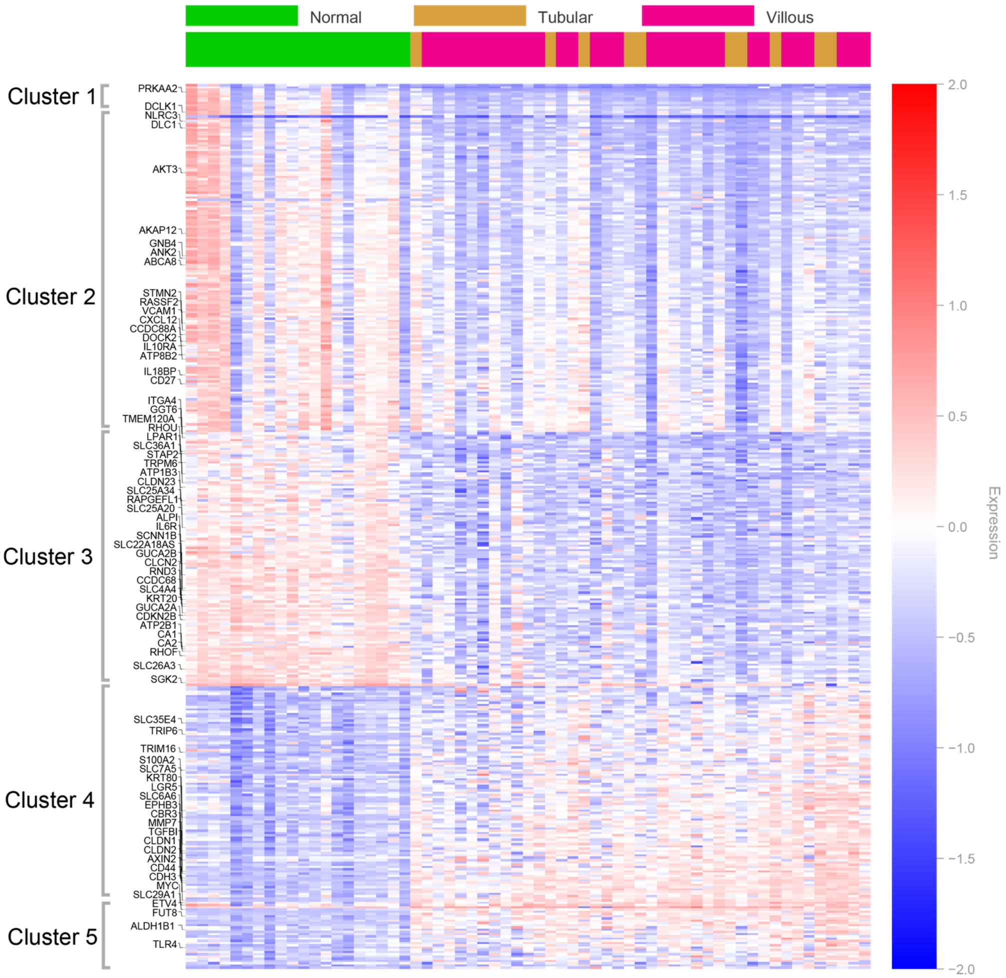

| | Precision [TP/(TP + FP)] | Recall [TP/(TP+FN)] | F1-score [2 x (precision x recall)/(precision + recall)] | Support [total samples] |
|---|---|---|---|---|
| **Normal (validation)** | 1.00 | 1.00 | 1.00 | 20 |
| **Adenoma (validation)** | 1.00 | 1.00 | 1.00 | 41 |

TP = True Positives; FP= False positives; FN = False negatives
**ROC AUC** (Receiver Operator Characteristic - Area under curve) = 1.00
**Classification Accuracy Score:** 1.00
**Fisher Exact p value**: 1.603e-16

**Supplementary Figure 4:** Heatmap of the gene expression values using genes present in the C#1-2-3-4-5 path in validation cohort #1 (accession: phs001384.v1.p1). Key genes in the clusters are presented on the left of the heatmap. The expression level ranged from -1 to +1; the expression below 0 indicated low expression and represented with blue color and the expression above 0 indicated high expression and represented with red color. Sample ranking using the C#1-2-3-4-5 path is shown in the bar plot above the heatmap.

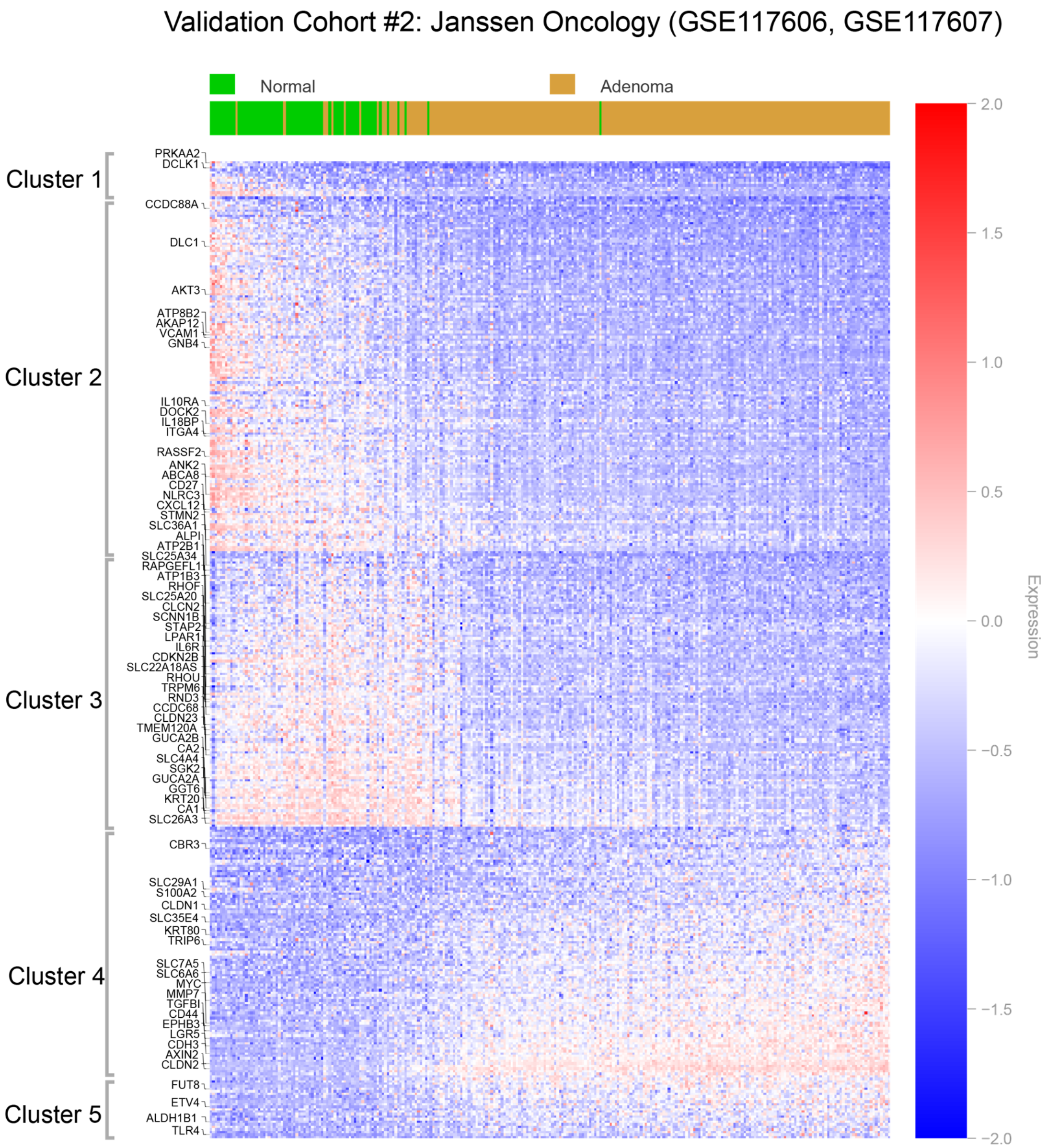

| | Precision [TP/(TP + FP)] | Recall [TP/(TP+FN)] | F1-score [2 x (precision x recall)/(precision + recall)] | Support [total samples] |
|---|---|---|---|---|
| **Normal (validation)** | 0.67 | 0.98 | 0.80 | 65 |
| **Adenoma (validation)** | 0.99 | 0.85 | 0.92 | 204 |

TP = True Positives; FP= False positives; FN = False negatives
**ROC AUC** (Receiver Operator Characteristic - Area under curve) = 0.98
**Classification Accuracy Score:** 0.88
**Fisher Exact p value**: 7.657e-37

**Supplementary Figure 5:** Heatmap of the gene expression values using genes present in the C#1-2-3-4-5 path in validation cohort #2 (GSE117606, GSE117607). Key genes in the clusters are presented on the left of the heatmap. The expression level ranged from -1 to +1; the expression below 0 indicated low expression and represented with blue color and the expression above 0 indicated high expression and represented with red color. Sample ranking using the C#1-2-3-4-5 path is shown in the bar plot above the heatmap.

Validation Cohort #3: PMID: 27270421 (GSE77953)

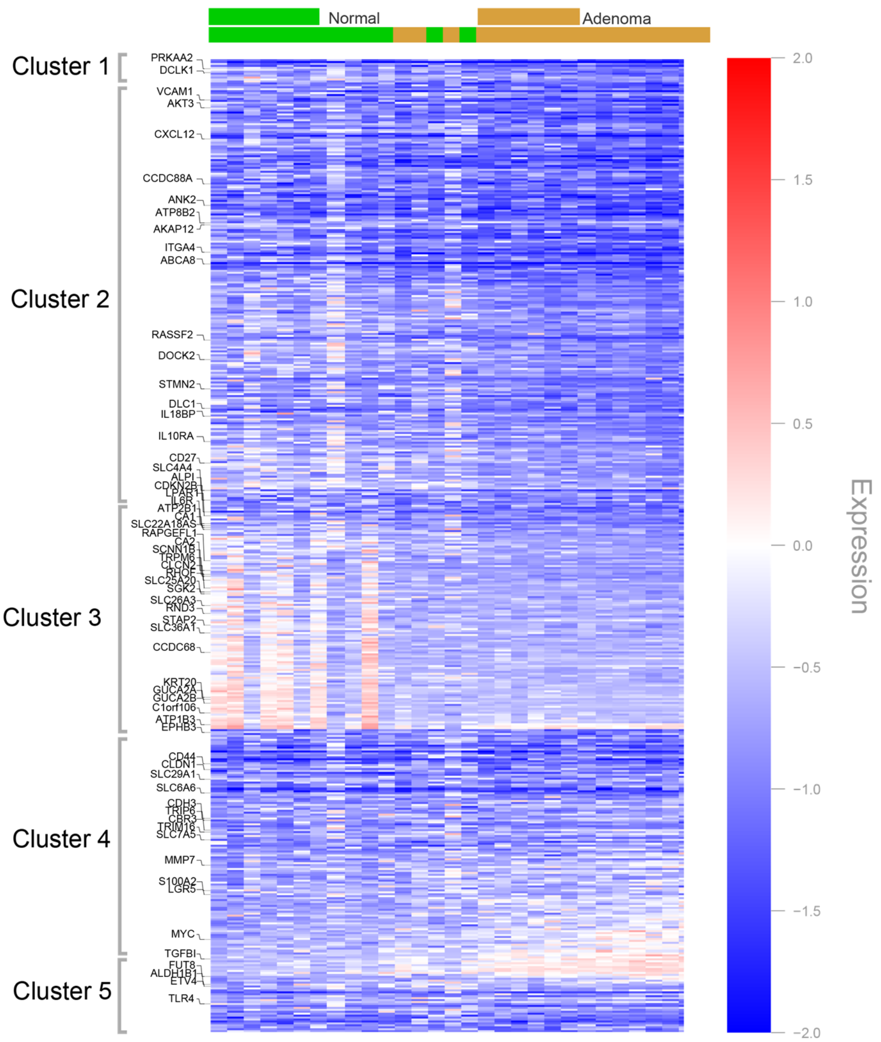

| | Precision [TP/(TP + FP)] | Recall [TP/(TP+FN)] | F1-score [2 x (precision x recall)/(precision + recall)] | Support [total samples] |
|---|---|---|---|---|
| **Normal (validation)** | 0.76 | 1.00 | 0.87 | 13 |
| **Adenoma (validation)** | 1.00 | 0.76 | 0.87 | 17 |

TP = True Positives; FP= False positives; FN = False negatives
**ROC AUC** (Receiver Operator Characteristic - Area under curve) = 0.98
**Classification Accuracy Score:** 0.866
**Fisher Exact p value**: 2.173e-05

**Supplementary Figure 6**: Heatmap of the gene expression values using genes present in the C#1-2-3-4-5 path in validation cohort #3 (GSE77953). Key genes in the clusters are presented on the left of the heatmap. The expression level ranged from -1 to +1; the expression below 0 indicated low expression and represented with blue color and the expression above 0 indicated high expression and represented with red color. Sample ranking using the C#1-2-3-4-5 path is shown in the bar plot above the heatmap.

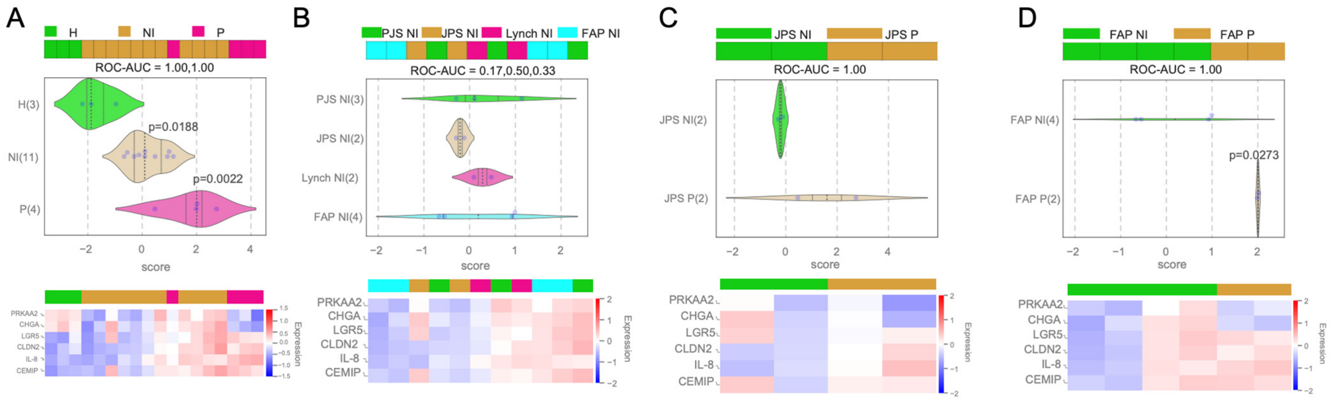

**Supplementary Figure 7**: Analysis of RT-qPCR on various samples based on DCT values. Genes included in the analysis are the core MACS genes: *PRKAA2*, *CHGA*, *LGR5*, *CLDN2*, *IL-8* and *CEMIP*. *Top*: Bar plot showing the separation of different samples based on the composite score. *Middle*: Violin plot showing the composite score for each of the samples by groups. The p-value from Student's t-test is shown if there is a significant difference ($p < 0.05$) between the first and selected group. *Bottom*: Heatmap of core MACS gene expression. **A)** Core MACS genes can separate healthy colon samples (n=3), colon samples from non-involved (NI) regions collected from different locations genetically predisposed patients (n=11) and colon samples from polyp (P) regions collected from genetically predisposed patients (n=4). **B)** Breakdown of colon samples from NI regions into specific diseases: PJS (n=3), JPS (n=2), Lynch (n=2) and FAP (n=4). **C)** Comparison of NI vs P regions from patients with JPS. **D)** Comparison of NI vs P regions from patients with FAP.

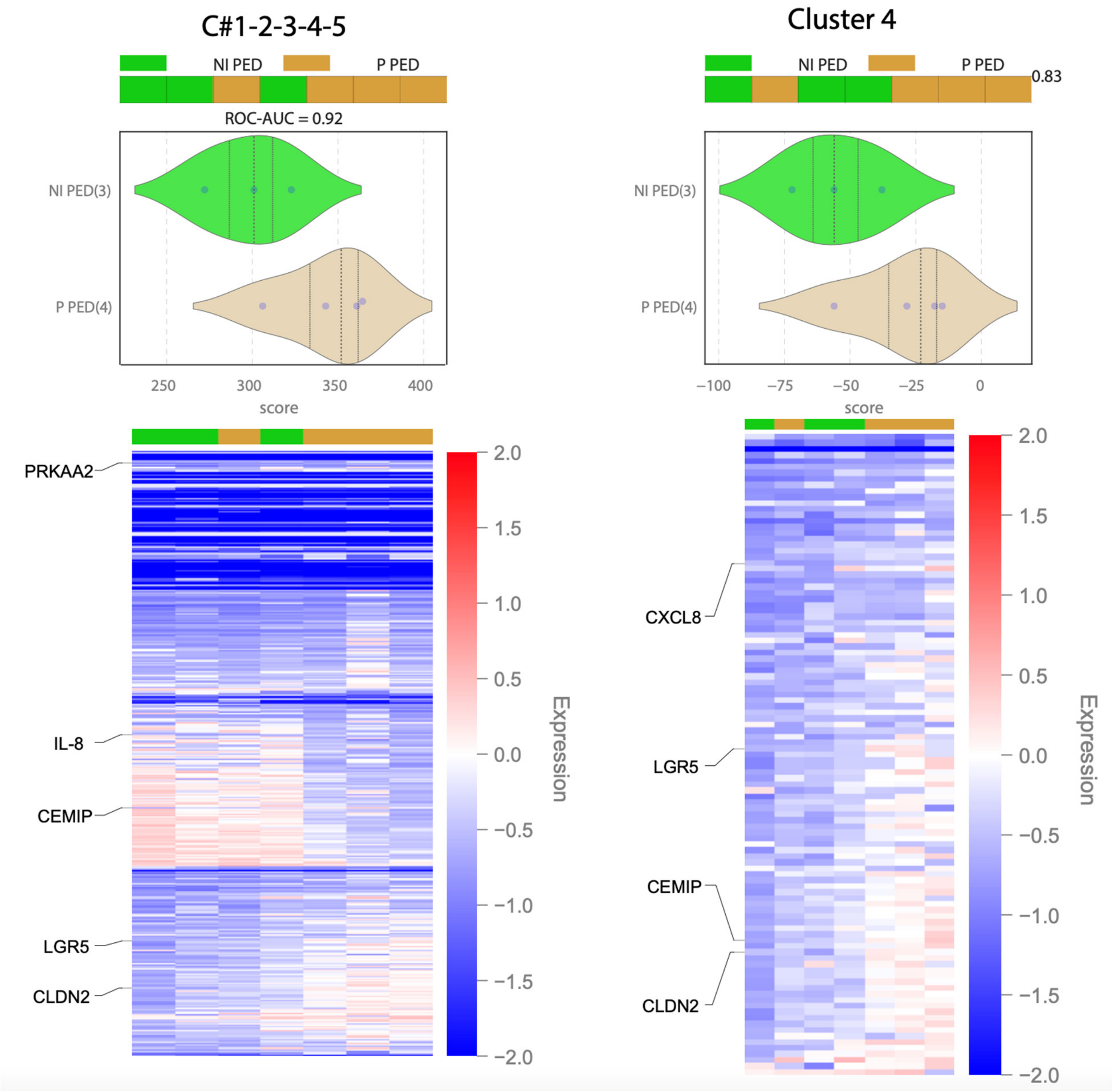

**Supplementary Figure 8**: *Top*: Bar plot and violin plot using the C#1-2-3-4-5 path (left) or C#4 (right) can segregate pediatric (PED) colon samples collected from non-involved (NI) regions (n=3) from colon samples collected from polyp (P) region (n=4). Samples were harvested from genetically predisposed CRC pediatric patients. Analysis of the gene expression was done by RNA-seq. *Bottom*: heat map showed the expression of the invariant genes from non-involved and polyp areas. Core MACS genes are presented on the left of the heatmap.

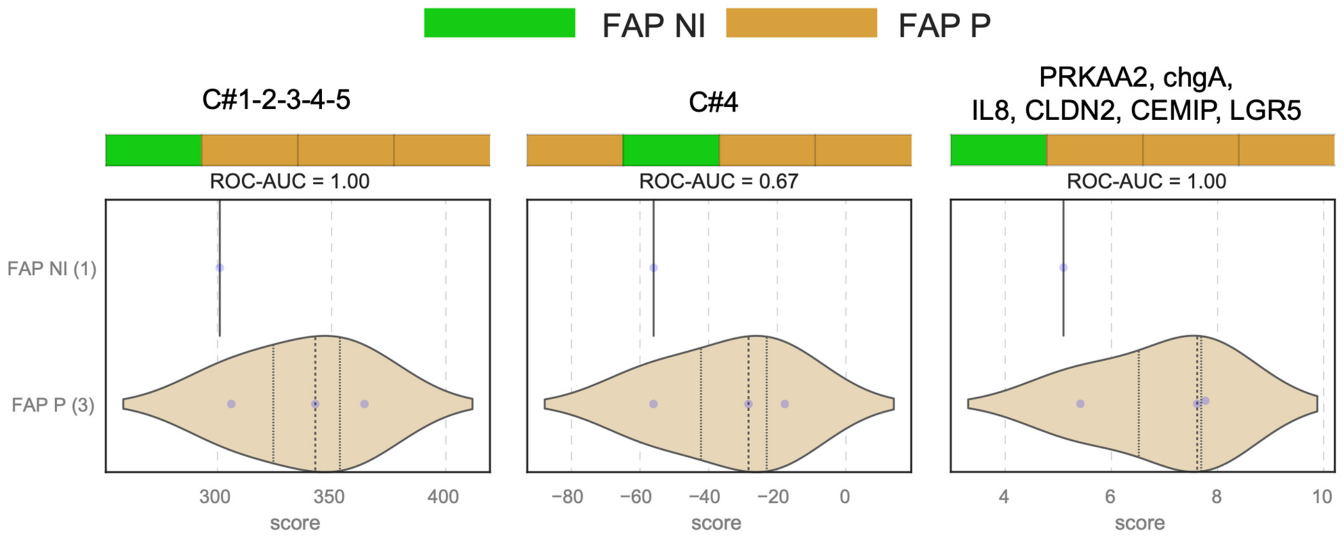

**Supplementary Figure 9:** Bar and violin plots using various gene signatures show separation of FAP non-involved (NI) from FAP polyp (P) samples that are obtained from patient derived organoids. Analysis of the samples was done by RNA-seq.

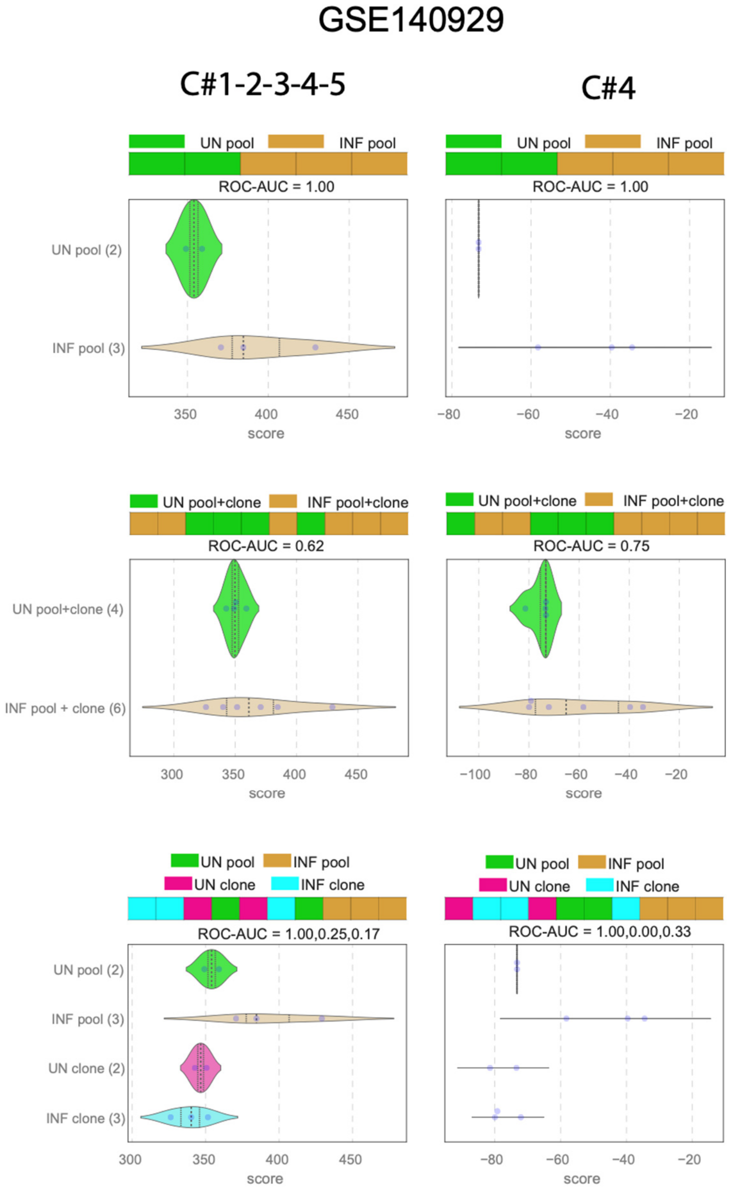

**Supplementary Figure 10:** *Top row:* Bar plots and violin plots show the separation of samples from uninfected organoids and organoids infected with pks+ *E. coli* (GSE140929). *Middle row*: Bar plots and violin plots shown the segregation of genes of uninfected primary mouse epithelial cells and epithelial cells infected with pks+ *E. coli*. *Bottom row:* Combined data of A & B. The sample ranks were based on genes in the C#1-2-3-4-5 path (left column) and C#4 genes (right column).

Boolean Pathway # 1-2-3-4-5 Can not separate uninfected compared to infected using probiotics and enteric pathogens

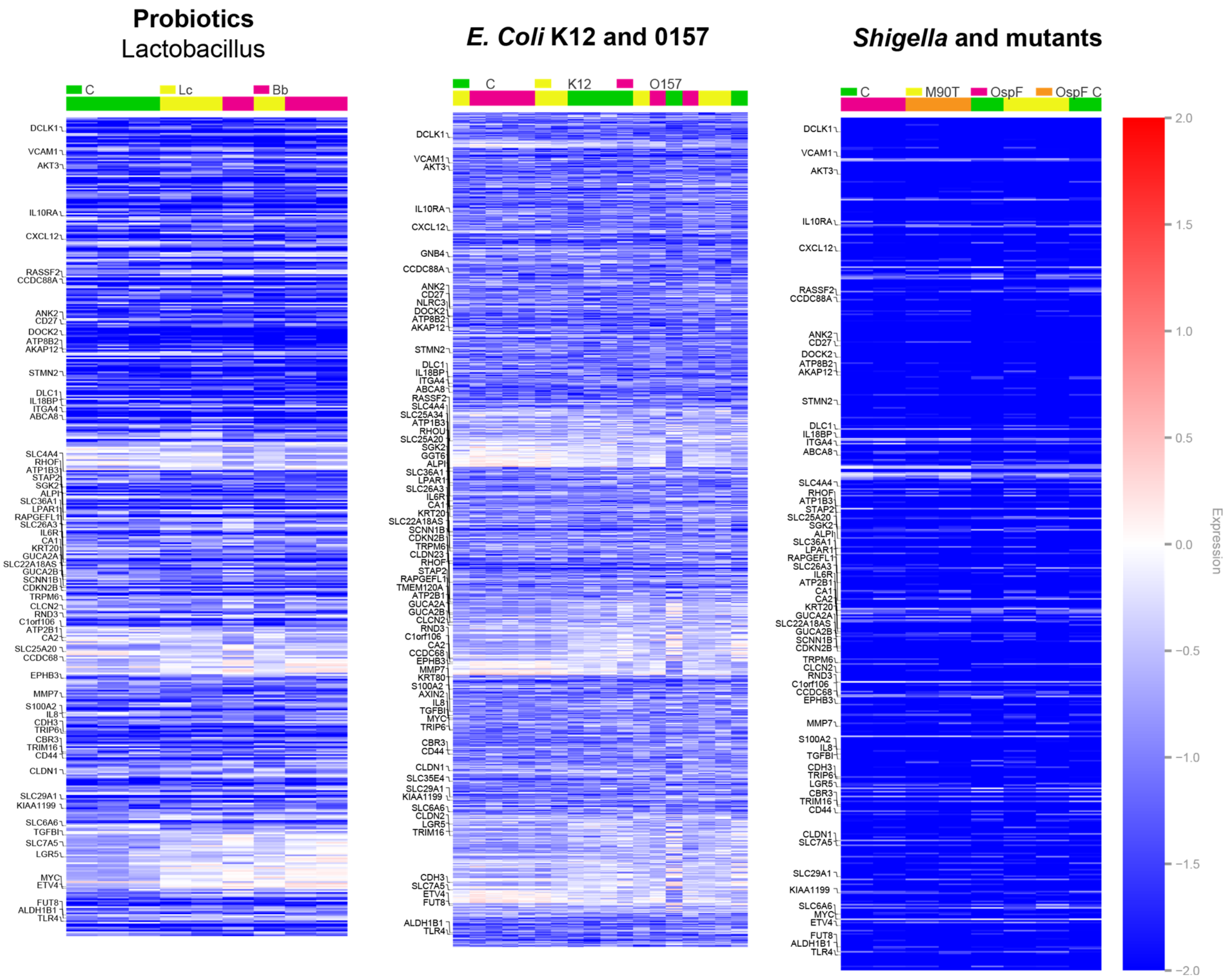

**Supplementary Figure 11:** The C#1-2-3-4-5 path cannot separate uninfected compared to infected using probiotics *Lactobacillus* (left) and enteric pathogens such as *E. coli* K12 and *E. coli* O157 strains (middle), and *Shigella* and mutants (right).
